# *Pseudomonas aeruginosa* CRISPR-Cas primes a minimal proline-codon toxin to abort anti-CRISPR phages

**DOI:** 10.64898/2026.06.02.729485

**Authors:** Rui Wang, Chao Liu, Ping Li, Jing Xu, Guangyi Liang, Yali Wu, Xinyu Jiang, Rui Zhao, Rafael Pinilla-Redondo, Shuai Le, Guangxin Luan, Ming Li

## Abstract

*Pseudomonas aeruginosa* typically carries type I-F CRISPR-Cas systems, which are targeted by diverse (pro)phage-encoded anti-CRISPR (Acr) proteins. Here, we show that ∼26% of *P. aeruginosa* Cas effectors are reprogrammed by a regulatory RNA guide (CreA) to transcriptionally silence a conserved small RNA toxin (CreT) that arrests cell growth upon Cas inactivation. This toxin unprecedentedly employs two consecutive proline codons, thus termed “proline-codon toxin”. Combined phylogenetic and genetic analyses unraveled that CreTA has forced *P. aeruginosa* to reject prophages whose Acrs disrupt Cas-DNA binding, which is essential for CreT repression. We further show that Acr-armed lytic phages designed to overcome CRISPR adaptive immunity can be instead aborted by CreTA-mediated defense *in vitro* and in a mouse model. Our findings unravel intra-genomic arms races between CreTAs and anti-CRISPR phages, and underscore the necessity of rational engineering of strain-specific therapeutic phages to penetrate the layered CRISPR and TA barriers of multidrug-resistant *P. aeruginosa*.

## INTRODUCTION

*Pseudomonas aeruginosa* is a Gram-negative opportunistic pathogen that frequently causes fatal infections in immunocompromised patients, including those suffering from cystic fibrosis, extensive burns, malignancy, or chronic obstructive pulmonary disease^1,2^. Its constitutive expression of multidrug efflux systems, low-permeability outer cell membrane, and chromosomally encoded antibiotic-modifying enzymes confers broad-spectrum intrinsic resistance that limits therapeutic options^3,4^. By re-directing the natural lytic activity of bacteriophages against multidrug resistant (MDR) *P. aeruginosa*^5–7^, phage therapy is attracting renewed interest as a next-generation antimicrobial strategy^8–11^.

Yet bacterial genomes have evolved diverse anti-phage defense mechanisms, including CRISPR-Cas, a widespread adaptive immune system that targets mobile genetic elements (MGEs) such as phages and conjugative plasmids^12–14^. Two classes, seven types (I-VII), and more than 30 subtypes have been defined^15,16^, with type I-F systems predominating in *P. aeruginosa*^17^. Each CRISPR locus contains short spacer memories of prior MGE exposure, interspersed with nearly identical direct repeats. CRISPR arrays are transcribed and processed into CRISPR RNAs (crRNAs) that direct Cas nucleases to cognate sequences, after which DNA cleavage occurs when the crRNA fully matches the invading protospacer and a flanking protospacer adjacent motif (PAM) is present^18,19^.

To neutralize this barrier, MGEs have evolved an anti-CRISPR (Acr) arsenal spanning >100 protein families^20,21^. These Acr proteins operate at multiple stages by blocking assembly of the Cas-crRNA ribonucleoprotein complex, disrupting the recognition of target DNA, or inhibiting the recruitment/activation of catalytic Cas components/domains. Notably, the first Acrs characterized were those that inhibit *P. aeruginosa* I-F systems^22^. More recently, RNA-based anti-CRISPRs (Racrs) that mimic crRNAs to sequester Cas proteins into non-functional complexes have also been discovered^23^. Because these Acr/Racr elements are widespread and highly diverse, they pose a serious threat to CRISPR-based immunity and simultaneously provide templates for designing therapeutic phages that can breach the CRISPR shield of MDR pathogens^24^.

Recent studies have uncovered a second, innate defensive layer in which CRISPR-Cas functions as a transcriptional regulator rather than a nuclease^25,26^. Many *cas* operons carry minimal, degenerate CRISPR arrays that produce crRNA-like RNAs (crlRNAs) ^27,28^. These noncanonical guides form only partial, PAM-proximal matches to endogenous promoter DNA, recruiting Cas effectors for repression instead of cleavage. The silenced targets are usually the *cas* operon itself^28^or small, cryptic toxin genes embedded within or adjacent to it^25,26,29^. In the latter case, the crlRNA regulator is termed CreA (CRISPR-resembling antitoxin) and the repressed toxin CreT (CRISPR-repressed toxin)^25^. When Acr proteins or Racr RNAs compromise the target DNA-binding activity of Cas proteins, CreT is de-repressed to induce dormancy or death, purging the Acr-encoding MGEs from the bacterial population^26^. This viral-effector-triggered innate immunity (independent of CRISPR targeting) is proposed to provide an “anti-anti-CRISPR” effect at the population level^30^. However, the influence of CreTA modules on the spread of Acr-encoding prophages and on the dynamics of lytic-phage infection remains unexplored.

Here we performed a systematic search for CreTA modules within type I-F CRISPR-Cas loci from diverse *P. aeruginosa* isolates and identified two conserved architectures, designated CreTA1 and CreTA2, in ∼26% of the loci. All these modules exploit a previously unrecognized yet conserved small RNA toxin featuring two consecutive proline codons—the most slowly decoded triplets in the genetic code. We therefore term it the “proline-codon toxin”. Combined phylogenetic and genetic analyses unravel an intense evolutionary arms race between these CreTA elements and prophage-encoded Acr proteins: only Acrs that do not titrate Cas away from the CreT promoter are tolerated and observed in the same genome. We further showed that lytic phages engineered to express CRISPR-inhibiting Acrs are instead neutralized by CreTA-mediated innate immunity. Our i*n vivo* infection data further underscore that therapeutic phages should be designed strain-by-strain to avoid the counter-productive outcome of Acr deployment.

## RESULTS

### Two conserved CreTA modules in *P. aeruginosa* I-F CRISPR-Cas loci

Most *P. aeruginosa* genomes carry a type I-F CRISPR-Cas cassette composed of *cas1*, a fused *cas2-3*, and a *csy* operon (*csy1-csy4*) encoding the Csy effector complex (Figure 1A). By manually exploring these loci, we identified two conserved crlRNA genes, almost always embedded in the intergenic region between *cas2-3* and *csy1* (Figure 1A and Table S1). This intergenic region is absent in the well-characterized PA14 I-F *cas* operon, likely explaining why these small RNAs went unnoticed. For each crlRNA gene, a nearby target site was predicted, flanked by the canonical I-F PAM (5′-CC-3′) (Figure 1A and Table S1). We also identified a small putative toxin gene downstream of each target site (see Figure 1A and below), resembling the architecture of CreTA systems and suggesting that these crlRNAs function as antitoxins. We therefore designate these crlRNAs as CreA1 and CreA2, and their cognate toxins as CreT1 and CreT2. CreA1 and CreA2 share similar repeat-like (ΨR) sequences, but possess different spacer-like (ΨS) guiding sequences: CreA1 favors a 20-nt ΨS, whereas CreA2 prefers a 26-nt ΨS (Figure 1A and Table S1).

**Figure 1.**
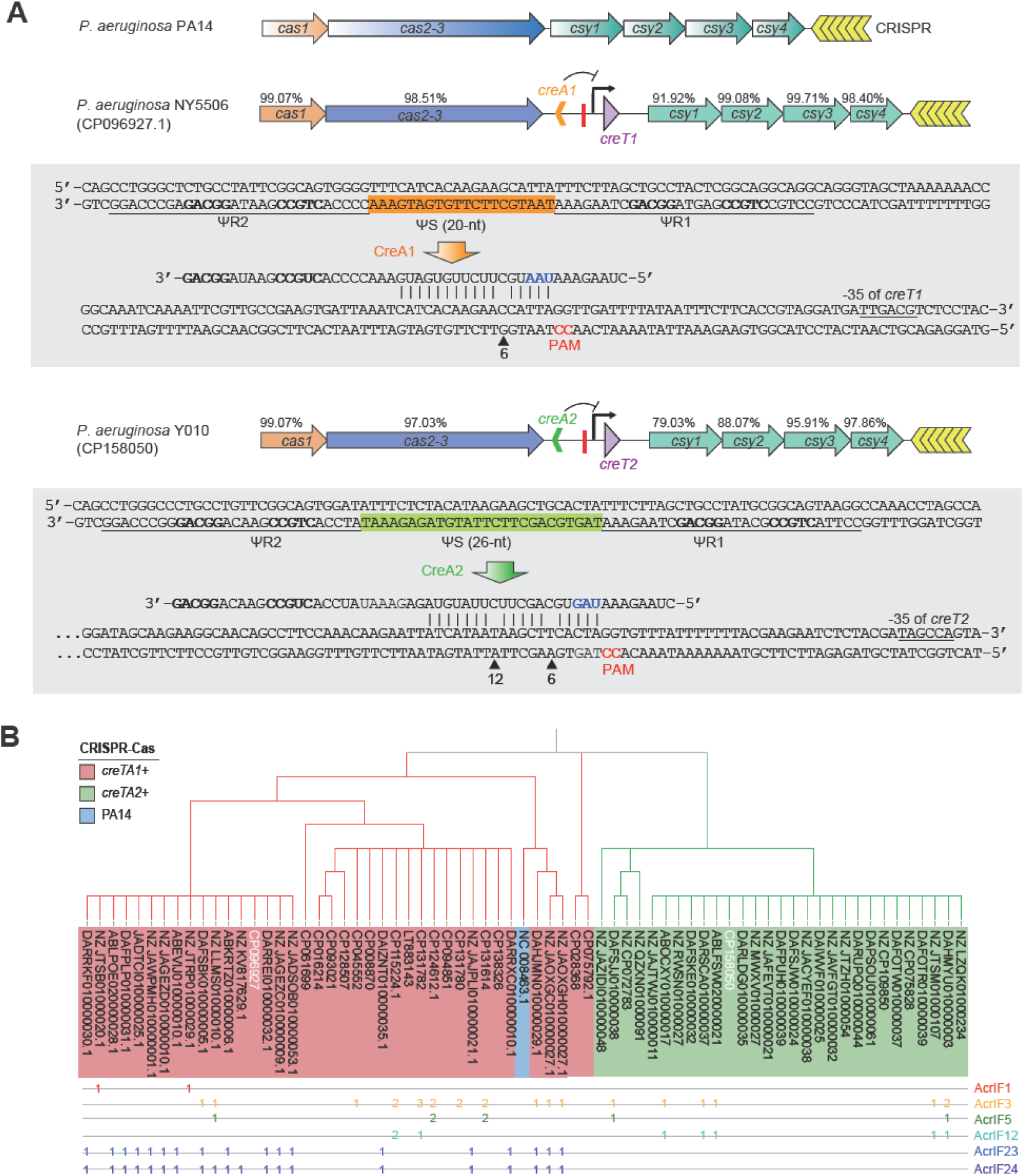
Two conserved CreTA modules embedded within *P. aeruginosa* type I-F CRISPR-Cas loci. (A) Architecture of representative loci carrying CreTA1 and CreTA2, respectively. Base pairings between CreA spacers (ØS) and their DNA targets are denoted by ‘|’, with PAM nucleotides highlighted in red. Mismatches at positions 6 and 12 (relative to the PAM) are marked. Blue nucleotides were subjected to mutational analyses in Figure 2. Palindromic nucleotides within the CreA repeat (ΨR) are bolded. Percent amino-acid identity of each Cas protein to its PA14 homolog is shown. (B) Maximum-likelihood tree of Csy1 sequences from CreTA-positive *P. aeruginosa* strains. Numbers indicate how many copies of a specific Acr protein are encoded in the same genome. The two strains depicted in (a) are highlighted in white; the PA14 type strain is shaded blue. See also Tables S1

To understand their evolutionary relationship, we constructed a Csy1-based phylogeny for these manually identified CreTA-positive *P. aeruginosa* genomes (Figure 1B). The tree resolves 40 genomes that encode CreTA1 and 29 that encode CreTA2 into two separate clades, indicating that the TA modules have experienced independent and parallel evolution. Because each clade contains both clinical and/or environmental isolates (Table S1), the association with CreTA modules seems not to be confined to any single ecological niche, but instead reflects broad selection pressures.

### CreA1 and CreA2 orthogonally regulate their cognate CreT toxins

We selected the CreTA1 module from strain NY5506 and the CreTA2 module from Y010 for functional characterization. Each *creA* contains two ΨR sequences that are less conserved than bona-fide CRISPR repeats (Figure S1A). Compared to ΨR2, ΨR1 retains the terminal 5′-CTAAGAAA-3′ motif that corresponds to the conserved 5′-handle on mature RNA guides. Nevertheless, both ΨR1 and ΨR2 retain the internal palindrome that forms a conserved RNA hairpin required for Csy4 processing (Figure S1B). Spacer-specific probes detected mature CreA1 and CreA2 RNAs (assumed to be 48-nt and 54-nt, respectively) in cells expressing the PA14 Cas machinery (Figure S1C), showing that these degenerate repeats remain able to be recognized and processed by this heterologous system.

The CreA1 and CreA2 ΨS sequences both form extensive base pairings with a target site upstream of the CreT toxin genes, particularly in the PAM-proximal region, except at positions 6 and 12 that do not participate in target DNA recognition^31,32^ (Figure 1A). To determine the transcription start sites (TSSs) of *creT1* and *creT2*, we fused their predicted promoters to *mCherry* and performed primer extension using an *mCherry*-specific primer (Figure S2). This defined canonical −10 and −35 elements for both promoters (Figure 2A). Notably, the −35 elements lie ∼30 bp downstream of the CreA target sites (Figure 2A), prompting us to test whether CreA-guided Csy complexes silence *creT* promoters by occluding RNA-polymerase loading.

**Figure 2.**
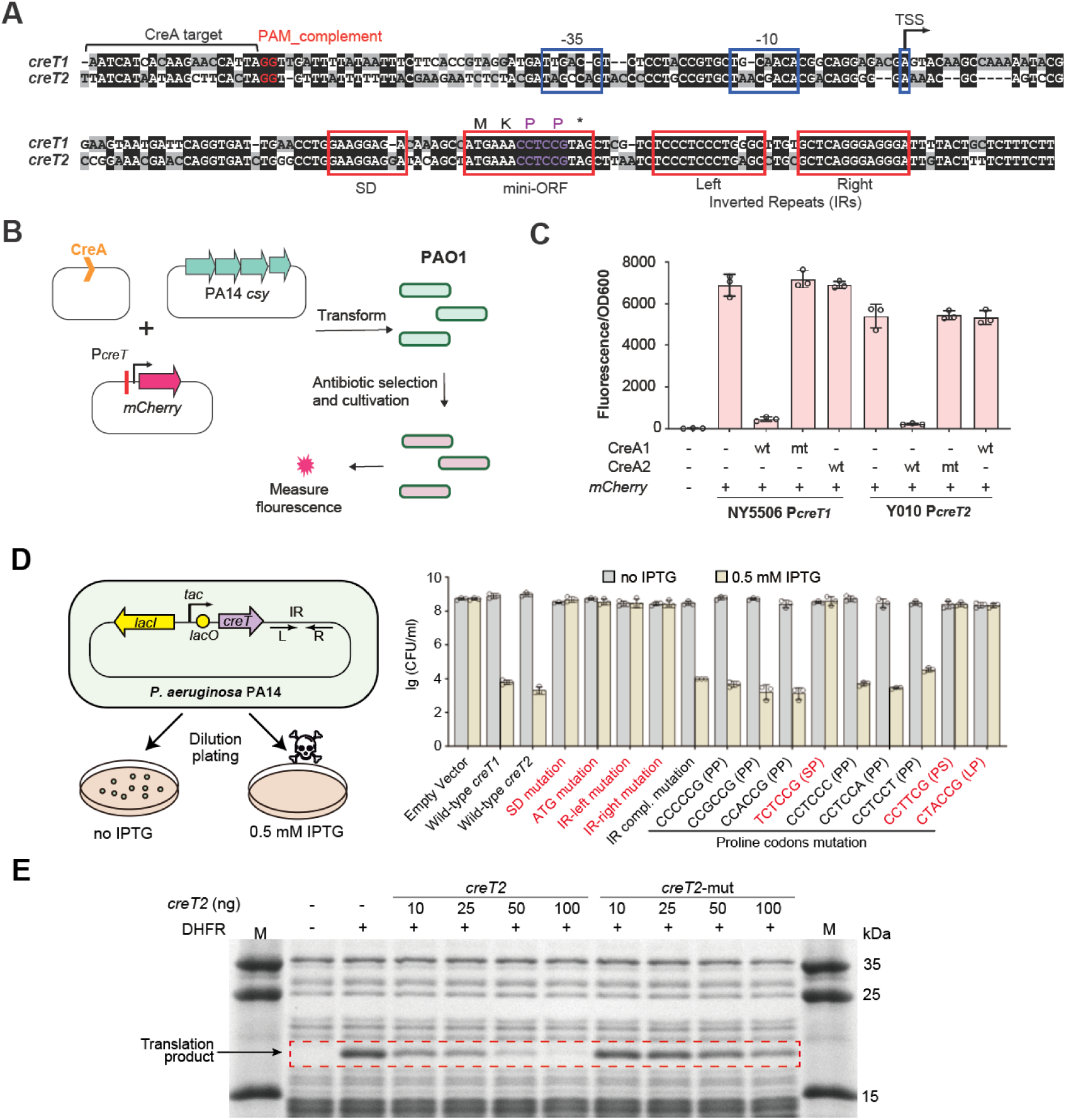
CreA1 and CreA2 transcriptionally silence a conserved proline-codon RNA toxin. (A) Sequence alignment of *creT1* and *creT2*. The promoter elements (−10 and −35) and the transcription start site (TSS) are blue framed. The conserved SD sequence, mini-ORF, and a pair of inverted repeats (IR), are red framed. The target site of cognate CreA and the complement of PAM are indicated. (B) Design of the fluorescence assay. (C) Fluorescence intensity from PAO1 cells with an *mCherry* gene under the control of P*_creT1_* or P*_creT2_* in the presence of PA14 Csy proteins. Three spacer nucleotides were mutated (shown blue in Fig. 1A) in CreA1 and CreA2 mutants (mt). wt, wild-type. Error bars, mean±s.d. (n=3). (D) Growth of PA14 cells expressing wild-type CreT1 or CreT2, or different CreT2 mutants (inactive mutants highlighted in red). The left (IR-left) and right (IR-right) portions of IRs were separately or complementarily mutated. Amino acid changes are indicated for proline (P) codon mutants (S, serine; L, leucine). (E) A coupled *in vitro* transcription-translation assay using DHFR production from a DNA template as the readout. T7 RNA polymerase was used. In *creT2*-mut, the two critical proline codons were mutated. M, protein markers in kilodaltons (kDs). See also Figures S1, S2, S3, S4 and S5.

We constructed *mCherry* reporters driven by P*_creT1_* and P*_creT2_* (Figure 2B). In PAO1 cells expressing the PA14 Csy proteins, CreA1 or CreA2 specifically reduced fluorescence from its cognate reporter by >90 % (Figure 2C). Mutating the spacer of either CreA RNA to disrupt complementarity with their cognate toxin promoters fully relieved repression (Figure 2C). Thus, *P. aeruginosa* CreA RNAs guide Csy proteins to inhibit CreT promoters via limited but essential complementarity, likely by blocking RNA polymerase access. Notably, the inhibitory effects are strictly orthogonal, as CreA1 does not inhibit P*_creT2_*, and CreA2 does not recognize P*_creT1_* (Figure 2C), supporting that the two paralogous CreT/CreA pairs have divergently evolved without cross-talk (Figure 1B).

### CreT is a minimal proline-codon toxin

We next dissected the molecular basis of CreT toxicity. *creT1* and *creT2* are highly similar in sequence and carry the same 15-nt “minimal ORF” (mini-ORF) flanked by a canonical Shine-Dalgarno (SD) motif and a GC-rich palindrome that folds into a stable hairpin (Figure 2A). To define the *cis*-elements required for toxicity, we constructed a series of truncated *creT2* genes, placed each fragment under the control of the IPTG-inducible *tac* promoter, and introduced the plasmids into *P. aeruginosa* PA14 (Figure S3A; Figure 2D). Bacteria were spotted in ten-fold dilutions on plates with or without IPTG, and viability was assessed. A > 5-log reduction in colony forming units (CFU) on inducing plates was observed only when the transcript retained all three components—SD, mini-ORF, and hairpin—in their native configuration (Figure S3B). Additionally, mutating any one element abolished this reduction (Figure 2D), demonstrating that each is necessary for toxin activity. A combination of the corresponding elements from CreT1 also induced toxicity (Figure 2D), confirming that the mechanism is conserved.

The *creT* architecture resembles previously described small RNA toxins that sequester rare transfer RNAs (tRNAs) via consecutive rare codons^25,33^. However, we did not find any rare codons within the mini-ORF. To find out how CreT RNAs exert toxicity, we introduced nucleotide insertions or deletions immediately after the ATG start codon of CreT2. Strikingly, toxicity persisted only when three nucleotides were deleted (Figure S3C), underscoring the importance of downstream codons, which consist of two consecutive proline codons (CCT and CCG) immediately followed by a stop codon (Figure 2A). Substituting either proline codon with synonymous or non-synonymous alternatives revealed that retaining two consecutive proline codons here is required for toxicity (Figure 2D). Previous studies have shown that proline incorporation is inherently slow during translation and consecutive proline codons can induce ribosomal stalling^34,35^. Consistently, in an *in vitro* transcription-translation assay, CreT2 dramatically reduced the translation products in a manner dependent on the two proline codons (Fig. 2E; Figure S4A). Accordingly, we propose that *P. aeruginosa* CreT toxins, designated ‘proline-codon toxins’, represent a previously undescribed class of small RNA toxins that exploit consecutive, highly translated proline codons to induce severe translational arrest, although the underlying mechanism remains to be further determined. Dilution plating assay post IPTG induction confirmed that such toxins are bacteriostatic, as continuous induction for up to 6 hours halted growth without reducing CFU relative to the vector control (Figure S4B).

We noted that complementary substitutions preserving the stem base-pairings of the GC-rich hairpin did not affect toxicity (Figure 2D), demonstrating that the structure rather than the primary sequence of this element is essential. Northern blots further showed that disrupting the hairpin structure reduced transcript cellular levels (Figure S5A), underscoring its role in maintaining CreT RNA stability. Furthermore, inserting a 15-nt random ‘spacing’ sequence between the mini-ORF and the hairpin did not affect toxicity (Figure S5B), hinting that the hairpin’s influence is likely confined to RNA protection rather than toxin function.

### CreTA influences Acr distribution across *P. aeruginosa* genomes

The logic that CreA directs Cas proteins to inhibit CreT toxins indicates that the viral Cas-inhibiting Acr proteins should unleash CreT toxicity. We previously showed that CreTA can expel Acr-bearing MGEs from a bacterial population by inducing cell dormancy or death^26^. Yet to what extent CreTA influences the natural Acr repertoire within a bacterial species has remained unclear. A total of 11 AcrIF proteins have been discovered from *P. aeruginosa* (pro)phages and plasmids, including AcrIF1-AcrIF7, AcrIF11, AcrIF12, AcrIF23, and AcrIF24^22,36–38^. Nevertheless, within the 69 manually identified CreTA+ *P. aeruginosa* genomes, we detected only six Acr proteins—AcrIF1, AcrIF3, AcrIF5, AcrIF12, AcrIF23, and AcrIF24 (hereafter “co-existing Acrs”)—resident in the same chromosomes (Figure 1B), supporting that CreTA may limit the spread of other Acr proteins.

To obtain a comprehensive view, we analyzed a collection of 11,828 *P. aeruginosa* genomes downloaded from the NCBI nucleotide database. By searching for Csy1 homologs, we identified 4,177 strains that carry the type I-F CRISPR-Cas system (Figure 3A). Within this CRISPR+ group, 1,052 genomes (25.19%) encoded the CreTA1 module, whereas only 38 genomes (0.91%) encoded CreTA2, bringing the total prevalence of CreTA elements among type I-F loci to 26.10 % (Figure 3A). A Csy1-based phylogeny revealed that the most majority of CreTA1+ isolates form a single, tightly linked terminal cluster with very short internal branches, whereas the CreTA2-positive genomes group together on a long, separate branch (Figure 3B). The topology indicates that both modules were acquired recently and independently, followed by rapid clonal expansion of the CreTA1-positive lineage.

**Figure 3.**
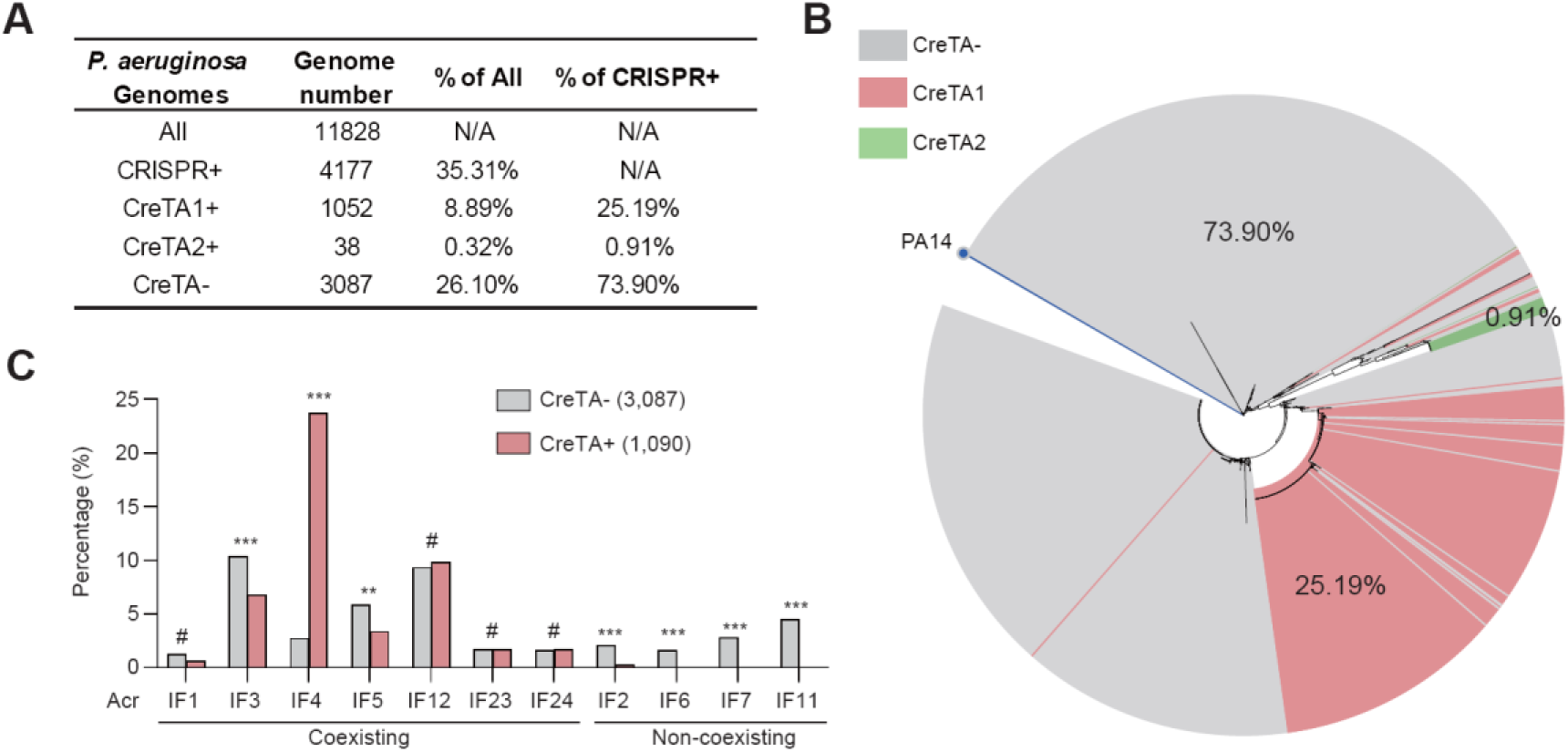
Distribution of CreTA and Acr elements across 11,828 *P. aeruginosa* genomes. (A) The number and percentage of *P. aeruginosa* genomes (downloaded from NCBI nucleotide database) carrying the type I-F CRISPR-Cas loci, with or without the embedded CreTA module. Percentages are also given for the fraction of CRISPR+ genomes that encode CreTA1, CreTA2, or lack CreTA. (B) Maximum-likelihood phylogeny of 4,177 *P. aeruginosa* Csy1 proteins. PA14 is marked for reference. (C) Percentage of CreTA- or CreTA+ genomes that carry a specific Acr protein. A two-tailed Fisher’s exact test was performed and *p-*values were adjusted for multiple comparisons using Benjamini-Hochberg false discovery rate (FDR) correction, #*p* > 0.05, \*\**p* < 0.01, \*\*\**p* < 0.001.

We next examined the distribution of Acr proteins across CRISPR+/CreTA+ (n = 1,090) and CRISPR+/CreTA-(n = 3,087) genomes (Figure 3C). AcrIF6, AcrIF7, and AcrIF11 were strictly confined to the CreTA-group, while AcrIF2, although occasionally detected in CreTA+ genomes, was strongly depleted from this cohort (*P* = 1.32 × 10⁻⁴, Fisher’s exact test). These findings indicate that these four Acrs are incompatible with CreTA and we therefore classify them as “non-coexisting.” In contrast, AcrIF1, AcrIF3, AcrIF4, AcrIF5, AcrIF12, AcrIF23, and AcrIF24 were frequently detected in both groups (AcrIF3 and AcrIF5 exhibited a measurable bias towards the CreTA-group), demonstrating that these Acr families can readily coexist with CreTA modules. Notably, AcrIF4, which is absent from our manually identified CreTA+ *P. aeruginosa* strains (Figure 1B), was present in 23.76 % (259/1090) of the bioinformatically defined CreTA+ genomes and displayed a significant positive association with this group, confirming its compatibility with CreTA (Figure 3C). Collectively, the data reveal that CreTA does influence Acr distribution across *P. aeruginosa* genomes.

### Non-coexisting Acrs induce CreTA toxicity

We then examined how these Acrs differently induce CreTA toxicity. We first constructed a chimeric operon, PA14-CreTA1, that combines the PA14 *cas* genes with the NY5506 intergenic region harboring *creTA1*, and introduced it into *P. aeruginosa* PAO1 (lacking native CRISPR-Cas apparatus) (Figure 4A). Each Acr was supplied on an arabinose-inducible PBAD plasmid. Strikingly, none of the ‘coexisting’ Acrs (AcrIF1, AcrIF3, AcrIF4, AcrIF5, AcrIF12, AcrIF23, and AcrIF24) reduced cell viability when expression was induced with arabinose, and CFUs were comparable to glucose-repressed controls (Figure 4B). AcrIF3, AcrIF4, AcrIF5, AcrIF12, and AcrIF23 evade toxicity because they block Cas2-3 recruitment, not the target DNA binding process^39–43^, thereby never allowing the toxin promoter to be derepressed. For AcrIF1 and AcrIF24, which instead disrupt the target DNA binding activity of Csy-crRNA complex^44,45^, we infer that an unknown mechanism shields the Csy-CreA complex from their action. Similarly, Acrs exhibiting coexistence with CreTA2 (AcrIF3, AcrIF5, and AcrIF12) neither induce this toxin module (Figure S6).

**Figure 4.**
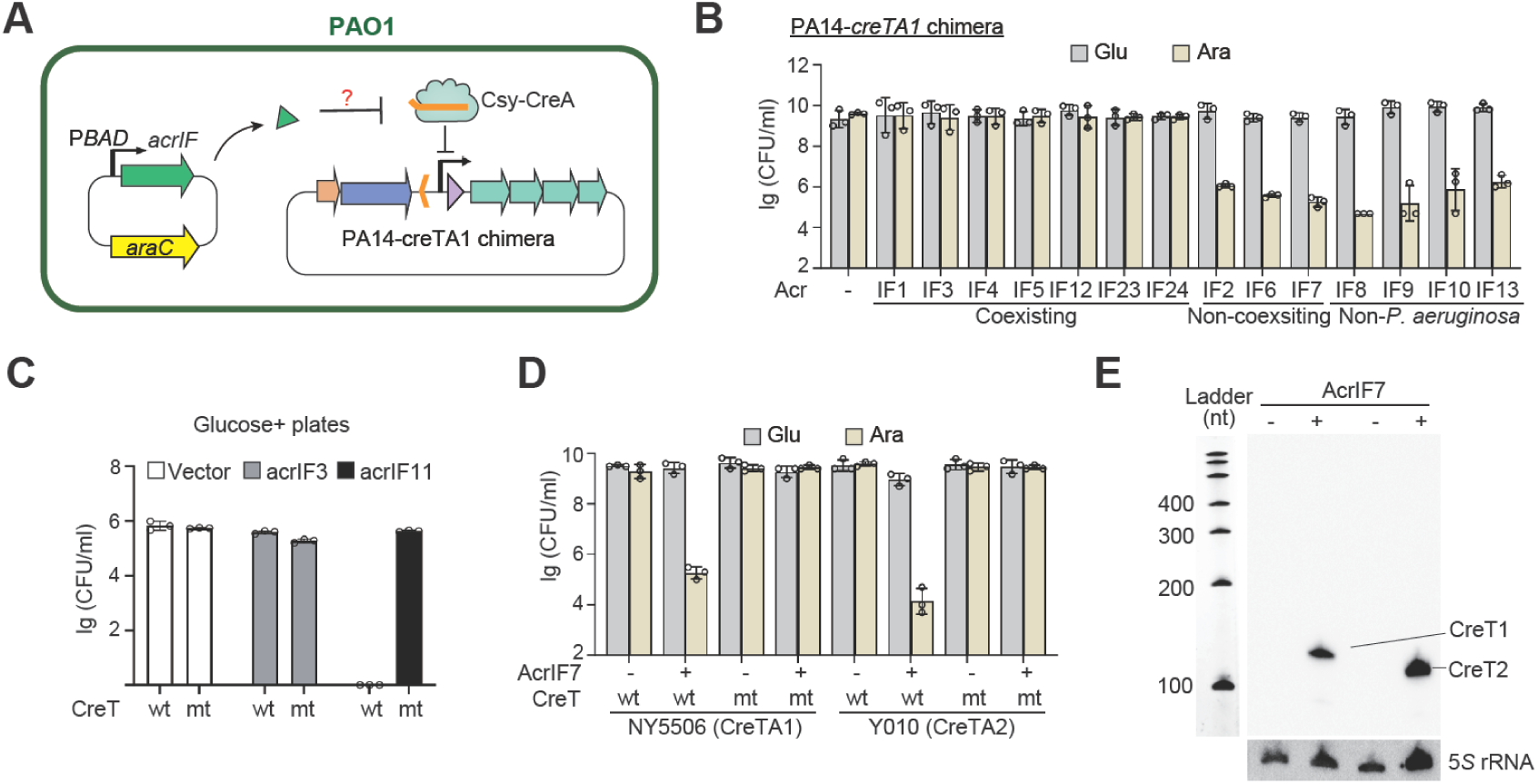
Differential activation of CreTA1 by Acr proteins. (A) Experimental design. *acr* genes were placed under the arabinose-inducible PBAD promoter. The NY5506 intergenic region encompassing *creTA1* was engineered into the PA14 *cas* operon to create a chimeric locus. (B) Growth of PAO1 carrying the PA14-CreTA1 chimera and an arabinose-inducible *acr* gene on medium supplemented with arabinose (Ara) or glucose (Glu). AcrIF proteins naturally cooccurring with CreTA1 in *P. aeruginosa* genomes are labeled “coexisting” (see Fig. 3c); the others are “non-coexisting” or “non-*P. aeruginosa*” (discovered in other bacterial species). (C) Transformation efficiency of PA14-CreTA1-containing PAO1 cells with plasmids encoding the indicated Acr proteins. A *creT*-mutated (proline codons deleted) strain served as a control. (D) Growth of PAO1 cells containing PA14-CreTA1 or PA14-CreTA2 together with a P*_BAD_*-controlled *acrIF7* on Ara- or Glu-containing medium. Error bars, mean±s.d. (n=3). (E) Northern blot detection of CreT1 and CreT2 transcripts in the presence (+) or absence (-) of AcrIF7 expression. 5*S* rRNA served as a loading control. A single-stranded DNA ladder was run in parallel. See also Figures S6.

In sharp contrast, AcrIF2, AcrIF6, and AcrIF7 tend to provoke the toxicity of CreTA1, causing a 3-5-log drop in CFU on arabinose plates (Figure 4B). The most efficient anti-CRISPR, AcrIF11, which catalyzes the ADP-ribosylation of the Csy complex^42^, could not be introduced into the strain containing PA14-CreTA1, even on plates that contain glucose inhibiting its expression (Figure 4C), indicating that its leaky expression is sufficient to trigger CreT toxicity. We further found that AcrIF8, AcrIF9, AcrIF10, and AcrIF13—discovered in species other than *P. aeruginosa*—likewise trigger CreTA1 toxicity (Figure 4B), implying that they would be counter-selected should they ever be horizontally transferred into CreTA1-bearing strains.

Specifically, we further showed that AcrIF7 induction reduced cell viability in the presence of wild-type CreTA1 or CreTA2, but not after we mutated their CreT toxins (Figure 4D), thus confirming that the observed toxicity derived from CreT rather than from the Acr protein itself. By Northern blotting, we further confirmed that CreT1 and CreT2 expression was induced by AcrIF7 (Figure 4E).

Together, these data reveal a striking compatibility rule: Acrs that naturally co-reside with a given CreTA module are unable to activate its toxin, whereas phylogenetically distant Acrs usually induce its toxicity. This suggests that the Acr-sensing innate defense line of CreTA forces bacterial genomes to reject those prophages (and likely plasmids) whose Acr proteins trigger the toxin, thereby manifesting in the exclusive distributions of CreTA and such Acrs across microbial communities.

### *P. aeruginosa* CreTA restricts conjugation by Acr-encoding plasmids

We then compared the efficacies of CRISPR-Cas-mediated targeting and CreTA-based innate defense in restricting conjugative plasmid transmission. A PAO1 derivative was engineered to carry either PA14-CreTA1 or PA14-CreTA2 together with a mini-CRISPR array, and conjugation frequencies were then measured for plasmids carrying a gentamicin-resistance cassette, the cognate protospacer, and/or an *acr* gene (Figure S7A).

When the protospacer was present, no gentamicin-resistant transconjugants were recovered (Figure S7B), confirming that CRISPR-Cas provides absolute, sequence-specific immunity. In the absence of the protospacer, the empty vector transferred at high frequency; however, plasmids expressing AcrIF7 or AcrIF11 still failed to yield resistant colonies (Figure S7B). Thus, CreTA-based innate immunity acts as an additional robust barrier against plasmid conjugation, thereby limiting the horizontal spread of antibiotic-resistance determinants.

### *P. aeruginosa* CreTA can abort phages engineered with Acrs

A recent study showed that engineered phages carrying *acr* genes efficiently kill multidrug-resistant *P. aeruginosa* despite being armed with CRISPR-Cas^24^. However, we asked whether CreTA can act as an “Acr sensor” and abort infections by such Acr-armed lytic phages.

To test this, we focused on investigating ΦYSTY4, a wild *P. aeruginosa* phage whose genome already contains *acrIF1* and the downstream *aca* regulator (Figure 5A). We deleted *acrIF1* while leaving *aca* intact (Δacr), or replaced *acrIF1* with *acrIF7* or *acrIF11* to create phages ΦIF7 and ΦIF11, respectively. Using PA14 cells with a plasmid producing a targeting crRNA, we showed that only the Δacr mutant lost the ability to evade CRISPR targeted immunity, whereas WT ΦYSTY4 and the ΦIF7 and ΦIF11 mutants all escaped surveillance (Figure S8).

**Figure 5.**
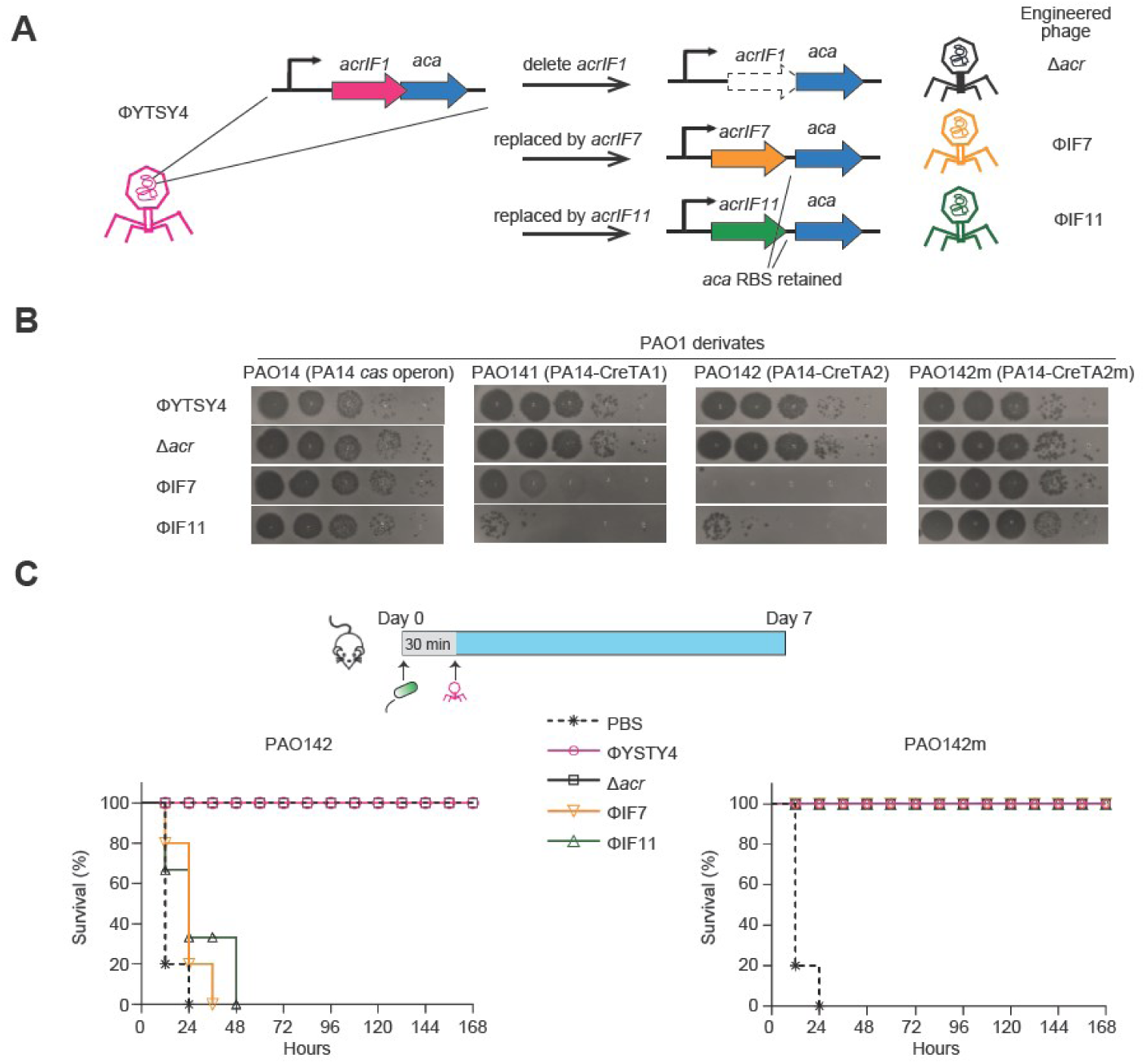
CreTA neutralizes lytic phages engineered with naturally ‘non-coexisting’ Acrs. (A) ÖYTSY4 engineering pipeline. Note that, because *acrIF1* and *aca* overlap, *aca* was kept intact during *acr* deletion and its ribosome-binding sequence (RBS) was deliberately retained upon replacement with *acrIF7* or *acrIF11*. (B) Plaque assays of wild-type (WT) or engineered ΦYTSY4 variants on PAO1 carrying: the PA14 *cas* operon (the strain denoted as PAO14), the PA14-CreTA1 chimera operon (PAO141), or the PA14-CreTA2 chimera (PAO142). For CreTA2, a control strain with a mutated *creT* (PAO142m) was included. (C) Therapeutic efficacy. 7-week-old BALB/c female mice were intraperitoneally inoculated with 100 μl *P. aeruginosa* (∼3 × 10^7^ CFU), followed 30 min later by 100 µl phage (∼3 × 10^8^ PFU). Each group included 5 mice. See also Figures S7 and S8.

These phages were then used to challenge PAO1 cells with a plasmid bearing the PA14 *cas* operon (denoted as PAO14), or the chimeric PA14-CreTA1 (PAO141) or PA14-CreTA2 (PAO142). In the absence of CreTA, these four phages showed equivalent titers in PAO1 cells (Figure 5B). However, when CreTA1 or CreTA2 was present, the titre of ΦIF7 and ΦIF11 dropped by >3-log compared with Δacr or WT ΦYSTY4, indicating that CreTA protects *P. aeruginosa* cells from these two Acr-armed phages. Consistently, the immune effect of CreTA2 vanished after we mutated its CreT toxin (Figure 5B). Notably, the WT phage encoding AcrIF1 maintained high titres on all PAO1 derivatives (Figure 5B), underscoring that deploying an Acr protein incapable of inducing CreTA is a reliable engineering strategy for increasing the effectiveness of therapeutic phages.

### Therapeutic efficacy of phages deploying Acrs

Finally, we tested the therapeutic performance of these phages using the mouse model. 7-week-old female BALB/c mice were first intraperitoneally challenged with ∼3 × 10⁷ CFU of *P. aeruginosa* PAO142 or PAO142m, and thirty minutes later, treated with ∼3 × 10⁸ PFU of ΦYTSY4, Δacr, ΦIF7, or ΦIF11 by the same route (Figure 5C). Notably, when the PAO142 infected mice were treated with a single dose of ΦIF7 or ΦIF11, all mice died within 2 days (Figure 5C). In contrast, a single dose of WT ΦYSTY4 or Δacr evaded CreTA-mediated anti-phage defense and rescued 100% of PAO142-infected mice. Nevertheless, in the PAO142m (CreT2 was mutated) infection group, all four phages 100% cured the infected mice (Figure 5C). Thus, arming phages with Acrs that trigger CreTA toxicity backfires when treating CreTA-positive *P. aeruginosa*.

Therefore, our *in vitro* and *in vivo* data demonstrated that CreTA can provide robust immunity against Acr-encoding lytic phages. This finding indicates that simply equipping a therapeutic phage with an *acr* gene may fail—or even backfire—if the pathogen carries a CreTA module instead of a targeting CRISPR spacer, and tailoring phages to deliver Acr proteins that silence CRISPR-Cas yet avoid inducing CreTA could provide a promising therapeutic avenue.

## DISCCUSION

Our study has uncovered cryptic ‘proline-codon toxins’ regulated by *P. aeruginosa* CRISPR-Cas systems. These toxins sense the activity of phage anti-CRISPR elements to induce cell death and abortive infection, revealing a deeper layer of the evolutionary arms race between *P. aeruginosa* and its invasive MGEs. CRISPR-repressed toxins (CreTs) were first described in haloarchaeal type I-B CRISPR systems, where small RNA toxins sequester rare arginine or isoleucine tRNAs^25,33^. More recently, a CRISPR-repressed small protein toxin was identified that safeguards type I-F systems in *Acinetobacter* species^26^. Proline-codon toxins now represent the third class of CRISPR-repressed toxins and appear to be specifically exploited by the type I-F systems of *P. aeruginosa*. It would be intriguing to explore why such toxins tend to be incorporated into *P. aeruginosa* CRISPR-Cas loci during evolution and whether they play additional roles in CRISPR-related or unrelated physiological processes. Nevertheless, the diversity of these CRISPR-regulated small RNA or protein toxins across different bacterial and archaeal species suggests that the integration of various small toxin genes into CRISPR-Cas loci may be a general paradigm, at least for type I CRISPR systems.

The discovery of *P. aeruginosa* CreTA modules, along with the well-characterized arsenal of anti-CRISPR elements in *P. aeruginosa* (pro)phages^22,36–38^, enabled us to examine, for the first time, the evolutionary conflict between CreTA modules and Acr elements. Our bioinformatic and experimental analyses revealed that Acr proteins encoded on the same chromosome as CreTA are unable to activate the CreT toxin, whereas Acr proteins lacking this linear coexistence are typically potent inducers of toxin expression and toxicity. In other words, CreTA negatively selects prophages or plasmids whose Acr effectors inhibit the DNA-binding activity of Cas proteins—the dominant mode of known Acrs^20,21^. Therefore, CreTA-based innate defense that senses a variety of Acr proteins constitutes an important barrier to Acr-encoding MGEs within bacterial communities, offering new insights into the arms race between host CRISPR systems and phage anti-CRISPRs.

Our bioinformatic analysis indicates that ∼26% of *P. aeruginosa* I-F CRISPR-Cas loci carry CreTA modules. Since prophages and conjugative plasmids often carry beneficial genes—such as antibiotic resistance determinants—CreTA may act as a barrier to the acquisition of these advantageous traits, potentially imposing evolutionary disadvantages on the bacterial host. We suppose the true prevalence of CreTA-positive CRISPR-Cas loci in *P. aeruginosa* is probably underestimated, considering that current databases are likely biased towards MDR isolates due to their clinical significance.

Moreover, CreTA-mediated innate immunity can abort the infection cycle of lytic phages. As demonstrated by our *in vitro* assays and murine infection model, engineered phages expressing Acr proteins can breach the adaptive defense provided by CRISPR-Cas but may inadvertently trigger CreTA toxicity, resulting in abortive infection. This finding underscores the importance of strain-specific customization in the selection or engineering of phage therapeutics, particularly given that the likelihood of a pathogen harboring a CreTA module is greater than the probability of it naturally carrying a CRISPR spacer targeting the therapeutic phage. It also suggests that engineering phages to deploy Acr proteins that block CRISPR-Cas but do not activate CreTA may be an effective therapeutic strategy.

In summary, we uncover the cryptic yet conserved proline-codon toxin embedded within *P. aeruginosa* CRISPR-Cas loci, underscoring the expanding repertoire of CRISPR-regulated toxins. Building on this discovery, we delineate the intra-genomic arms race between defensive CreTA modules and invasive Acr effectors, and provide practical guidelines for engineering phage therapeutics that evade both layers of immunity.

## RESOURCE AVAILABILITY

### Lead contact

Further information and requests for resources and reagents should be directed to and will be fulfilled by the Lead Contact, Ming Li.

### Materials availability

Materials are available from Ming Li upon request.

### Data and code availability

This paper does not report original code.

All data and materials reported in this paper will be shared by the lead contact upon request. This paper does not report original code.

## ACKNOWLEDGMENTS

We acknowledge Prof. Dongsheng Zhou (State Key Laboratory of Pathogen and Biosecurity, Beijing Institute of Microbiology and Epidemiology, Academy of Military Medical Sciences, Beijing, China) for generously providing the host strain of ΦYTSY4. We acknowledge Prof. Chengyuan Wang (Shanghai Institute of Immunity and Infection, Chinese Academy of Sciences) for helpful discussion on CreT toxicity mechanism. This work was supported by the National Key Research and Development Program of China [2024YFA0919400], the Strategic Priority Research Program of the Chinese Academy of Sciences [XDB0810000], the National Natural Science Foundation of China [32150020, 32230061, 32270092, 32370090, and 32200057], and the Youth Innovation Promotion Association of CAS [2020090].

## AUTHOR CONTRIBUTIONS

M. Li. conceived and supervised the project with valuable suggestions from S. Le. M. Li., G. Luan., S. Le., C. Liu. and R. Wang. designed the experiments. J. Xu., G. Liang., and C. Liu. performed the bioinformatics analyses. R. Wang., C. Liu., P. Li., Y. Wu. and J. Xu. conducted the efficiency of plaque assays. C. Liu. and Y. Wu. performed the fluorescence measurement. C. Liu. and R. Wang. performed primer extension assays. R. Wang. and C. Liu. performed Northern blot analysis. Y. Wu. and C. Liu. edited the phage genome. C. Liu., Y. Wu. and R. Wang. performed the phage infection assay. S. Le., X. Jiang. and R. Zhao. performed the animal infection experiments. M. Li., G. Luan., S. Le., C. Liu., and R. Wang. analyzed data. M. Li. prepared the original draft. R.P.-R. proofread the manuscript and offered valuable suggestions. M. Li., G. Luan. and S. Le. finalized the manuscript with input from all authors.

## STAR* METHODS

Detailed methods are provided in the online version of this paper and include the following:

- KEY RESOURCES TABLE
- RESOURCE AVAILABILITY

- Lead contact
- Materials availability
- Data and code availability
- EXPERIMENTAL MODEL AND STUDY PARTICIPANT DETAILS

- Bacterial strains and growth conditions
- METHOD DETAILS

- Phage purification
- DNA isolation and manipulation
- Bioinformatics analysis and phylogenetic tree construction
- In Vitro Protein Synthesis
- Phage infection assay
- CRISPR-Cas9 editing of the ΦYTSY4 phage genome
- Plasmid conjugation
- Fluorescence measurement
- RNA isolation
- Primer extension analysis
- Northern blot analysis
- Dilution plating assay
- Animals and Ethics Statement
- Mouse infection and phage therapy experiment
- QUANTIFICATION AND STATISTICAL ANALYSIS

## SUPPLEMENTAL INFORMATION

Supplemental information can be found online.

## STAR* METHODS

## KEY RESOURCES TABLE

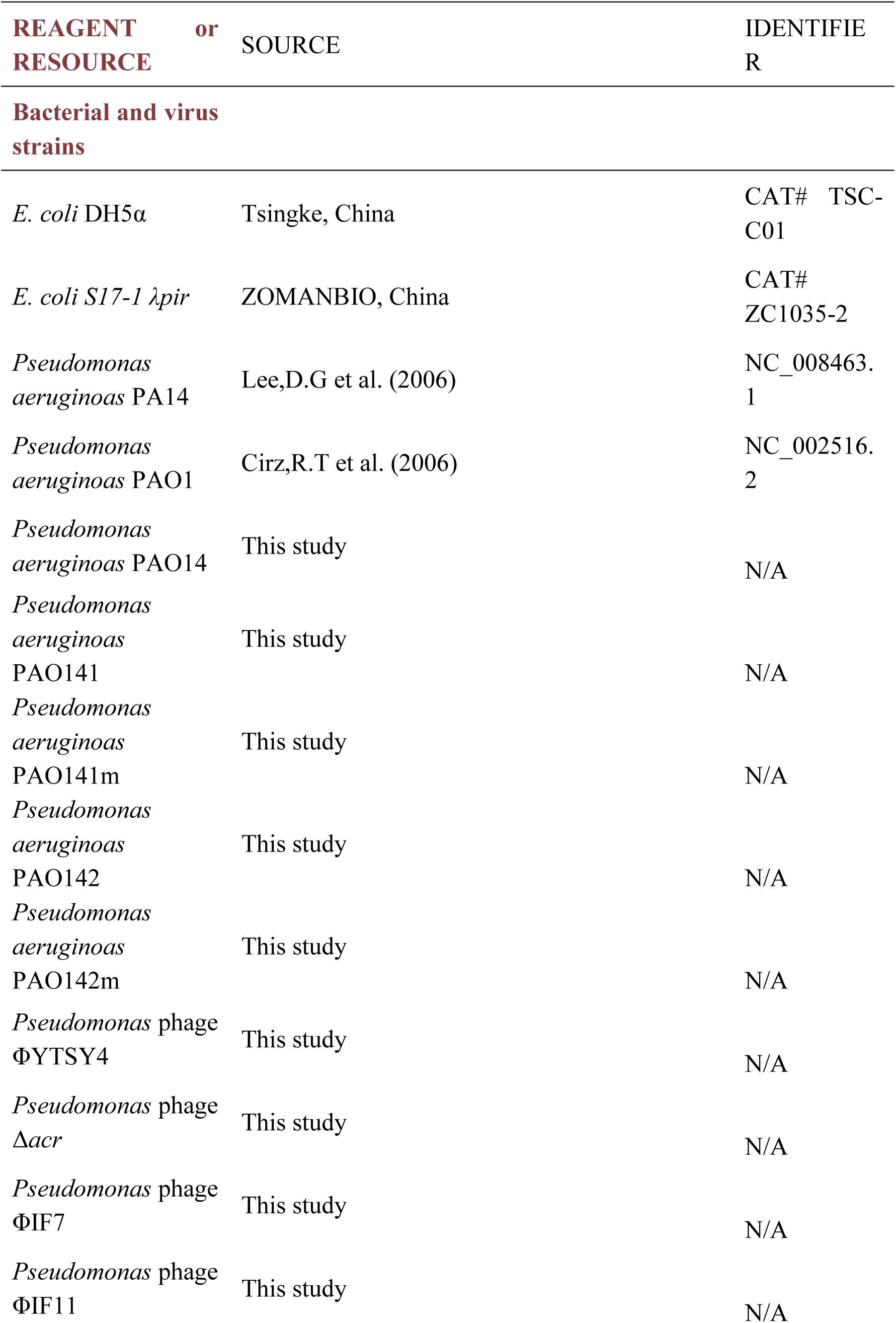
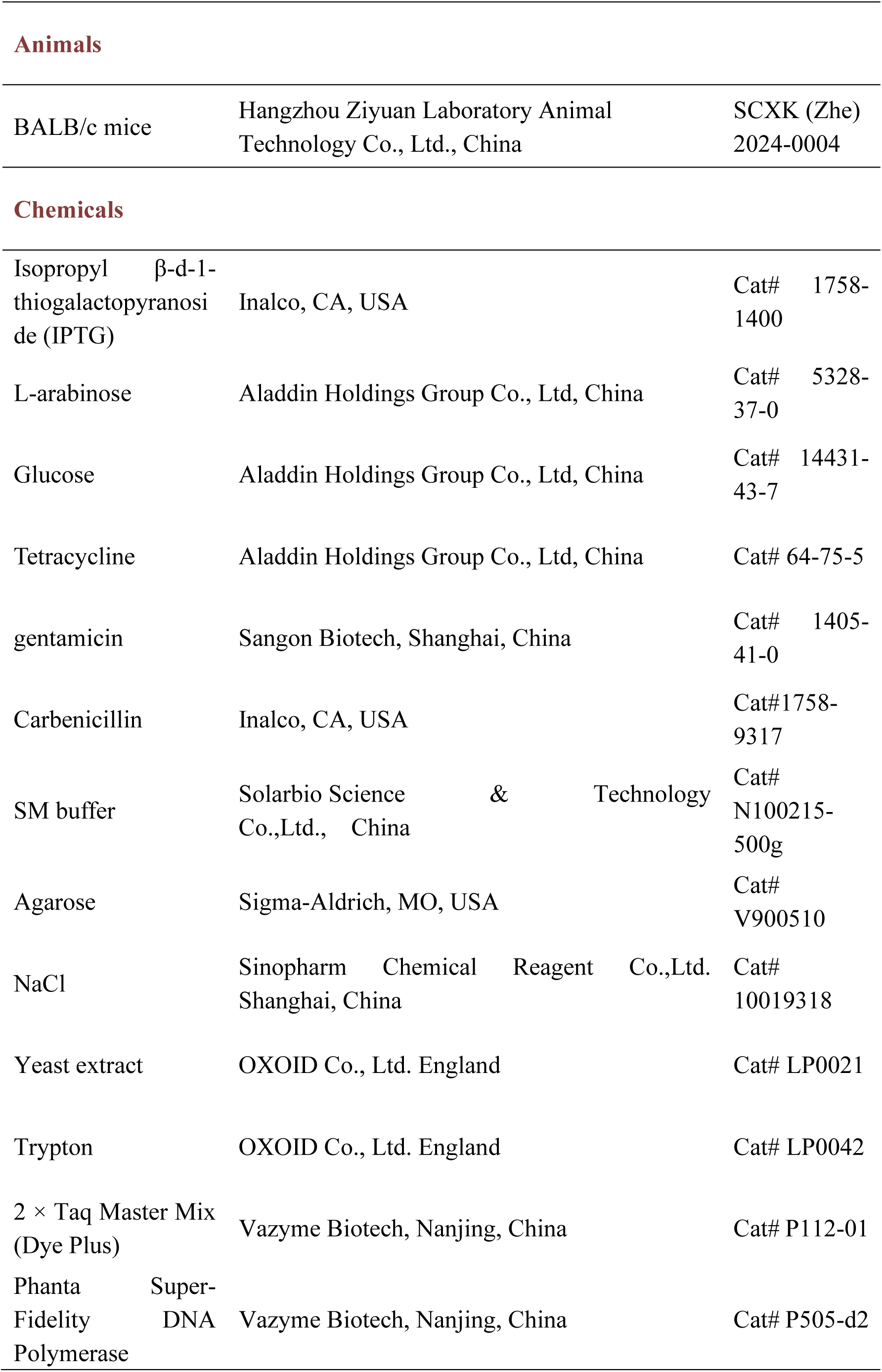
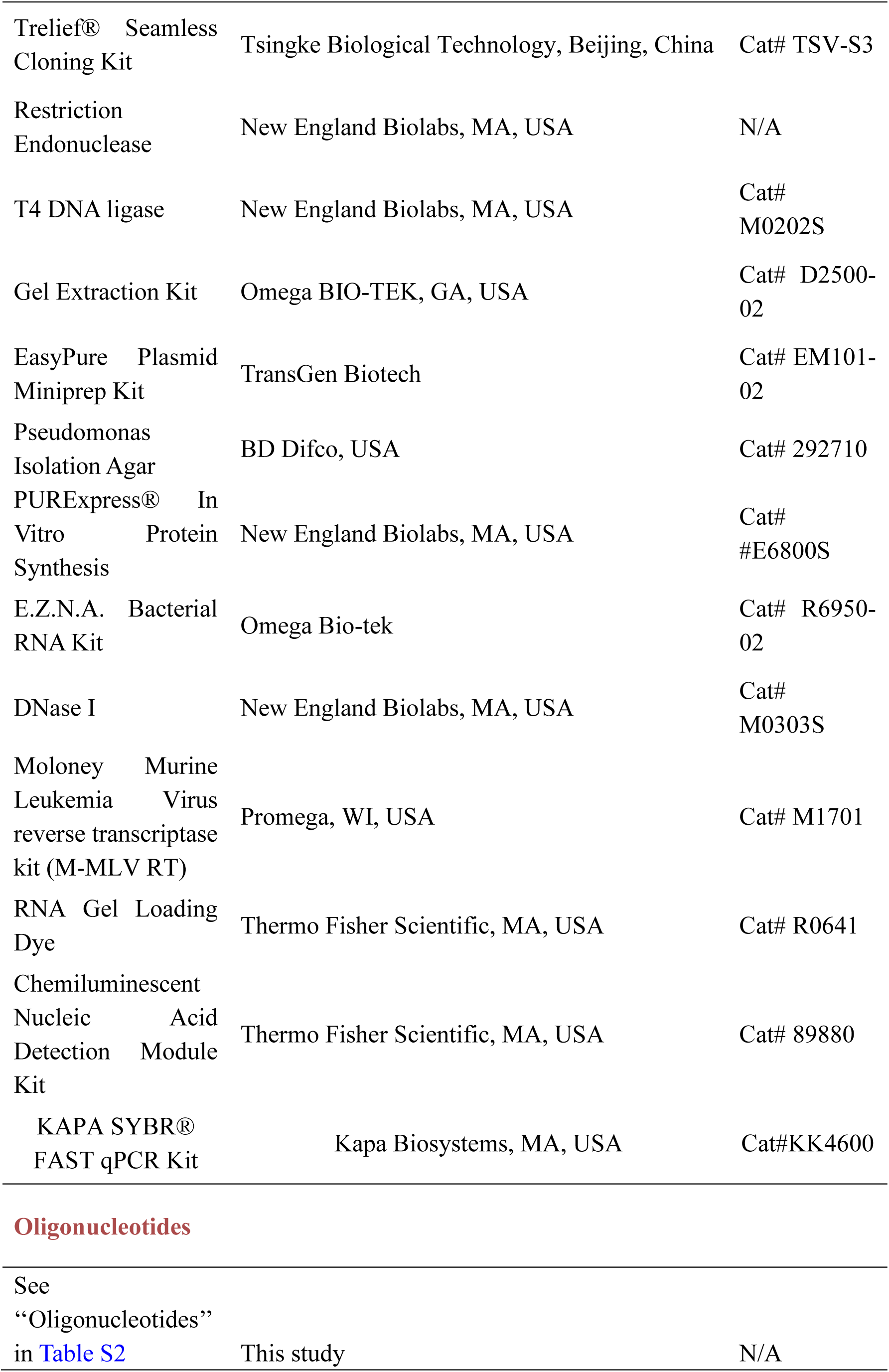
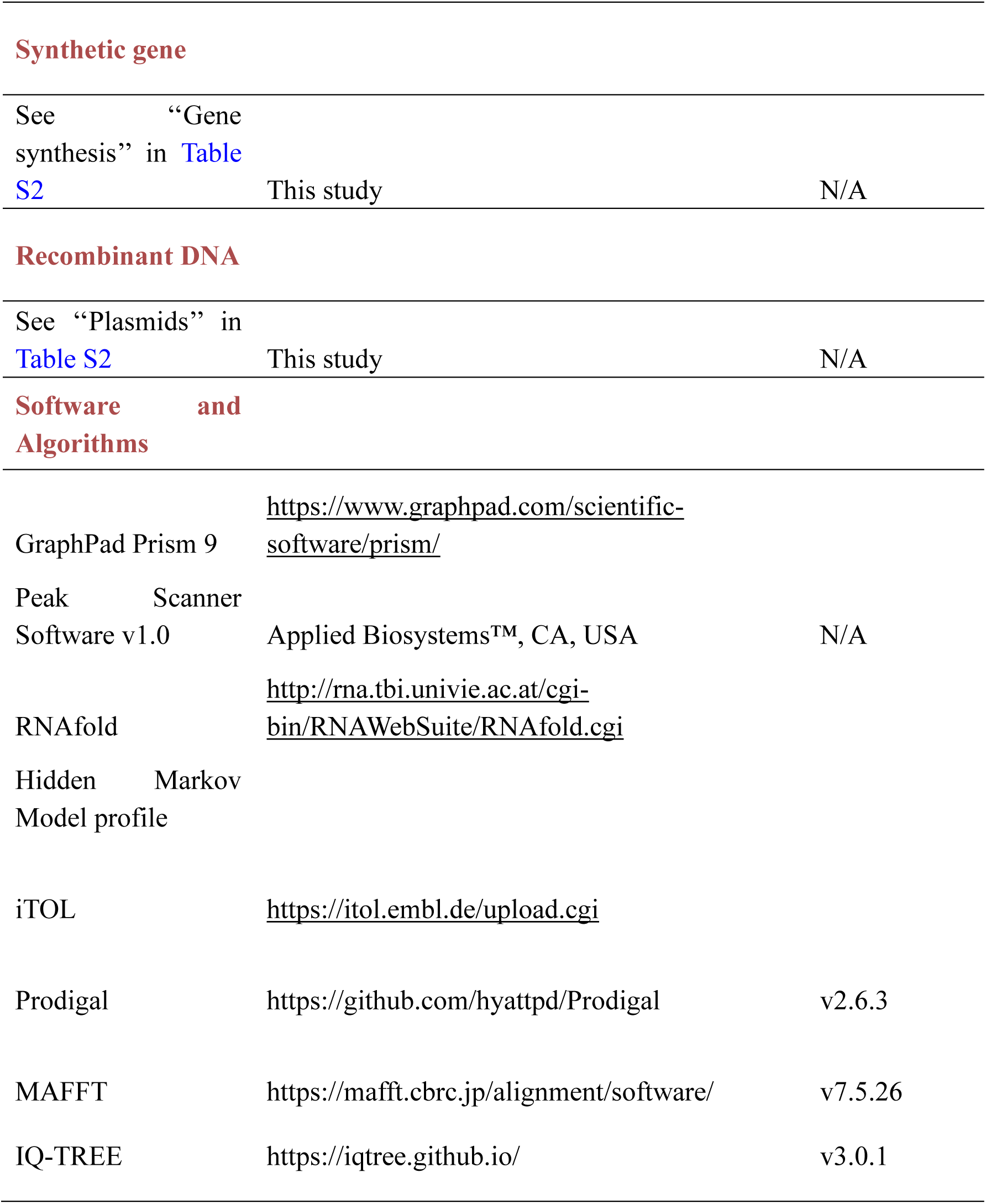

## EXPERIMENTAL MODEL AND STUDY PARTICIPANT DETAILS

### Bacterial strains and growth conditions

The bacterial strains used in this study included *Pseudomonas aeruginosa* PA14 and PAO1, as well as *Escherichia coli* DH5α and S17-1 λpir. All strains were routinely cultured in Luria-Bertani (LB) broth at 37℃ with shaking at 200 rpm. When required, antibiotics were supplemented at the following concentrations: for *P. aeruginosa*, carbenicillin (300 mg L⁻¹), gentamicin (50 mg L⁻¹), and tetracycline (50 mg L⁻¹); for *E. coli*, carbenicillin (100 mg L⁻¹), gentamicin (15 mg L⁻¹) and tetracycline (25 mg L⁻¹). Inducers and repressors, including isopropyl β-D-1-thiogalactopyranoside (IPTG, 500 μM), L-arabinose (1.0%, w/v), and glucose (0.5%, w/v), were used as specified in individual experiments.

## METHOD DETAILS

### Phage purification

To amplify ΦYTSY4, 1 mL of an overnight culture of the host strain *P. aeruginosa* PAO1 was inoculated into 100 mL of fresh LB medium and cultivated for one hour. A single plaque was then transferred to this culture and co-incubated overnight. Bacterial cells were removed by centrifugation at 4,000 × g for 10 min at 4 ℃, and the supernatant was filter-sterilized using a 0.22 µm filter. The phage titer was determined by spotting 2 µL of serial ten-fold dilutions onto a bacterial lawn prepared with 200 µL of an overnight host culture in a soft LB agar (0.5% w/v) overlay. After overnight incubation, plaques were counted to calculate the titer. The amplified phage was stored in SM buffer (100 mM CaCl_2_, 8 mM MgSO_4_, 50 mM Tris-HCl pH 7.5, and 0.01% w/v gelatin).

### DNA isolation and manipulation

All oligonucleotides used in this study are listed in Table S2. DNA polymerases, restriction enzymes, and T4 DNA ligase, as well as the Gibson Assembly Master Mix, were procured from Vazyme, TSINGKE or New England Biolabs (NEB). Standard protocols were followed for PCR, restriction digestion, Gibson assembly, ligation, and *E. coli* transformation. PCR products were purified using the E.Z.N.A. Gel Extraction Kit (Omega Bio-tek), and plasmid DNA was extracted from overnight *E. coli* cultures using the EasyPure Plasmid Miniprep Kit (TransGen Biotech). All constructed plasmids were verified by Sanger sequencing.

### Bioinformatics analysis and phylogenetic tree construction

A total of 11,828 *P. aeruginosa* genomes were obtained from the NCBI nucleotide collection. All genomes were uniformly annotated using Prodigal (v2.6.3) to predict protein-coding sequences. The *creTA* genetic elements were identified by performing a BLASTN search (identity ≥ 90%, query coverage ≥ 90%) against the genome database using known *creTA* sequences as queries. Candidate hits were manually curated to confirm their genomic context and distribution. Similarly, AcrⅠF protein homologs were identified from the predicted proteomes using hmmsearch (e-value ≤ 0.001) with a Hidden Markov Model profile constructed from known AcrⅠF protein sequences (https://docs.google.com/spreadsheets/d/17BR_Cy2jmokSKVZQwq7U6QgMUCUM5OBjnVnWbmsc6Eg/edit?pli=1&gid=361557639#gid=361557639), and their distribution across strains was assessed.

To construct the Csy1-based phylogeny, Csy1 protein homologs were identified from the annotated proteomes using hmmsearch (e-value ≤ 0.001) with a custom HMM profile built from 71 reference Csy1 sequences. The resulting Csy1 sequences were aligned with MAFFT (v7.5.26), and a maximum-likelihood phylogenetic tree was constructed using IQ-TREE (v3.0.1), which also determined the best-fit substitution model. The Csy1 homologs associated with CreTA1 or CreTA2 were mapped onto the tree using the iTOL (https://itol.embl.de/upload.cgi) platform for visualization.

### *In vitro* transcription-translation assay

T7 promoter and RBS were first linked to *creT2* or *creT2-mut* (two proline codons were mutated) by bridge PCR using synthetic primers, followed by purification with Gel Extraction Kit (Omega BIO-TEK). *In vitro* transcription-translation assay was then performed using the PURExpress In Vitro Protein Synthesis Kit (NEB), according to the manufacture’s instructions. Reactions were assembled in nuclease-free 0.5 mL microcentrifuge tubes, with a final volume of 5 µL per reaction. Each reaction contained 50 ng of DHFR control plasmid and, where indicated, 10, 25, 50, or 100 ng of the synthetic *creT* DNA constructs. After gentle mixing, the components were collected by brief centrifugation. Reactions were incubated at 37℃ for 2 hours and then placed on ice to terminate. For analysis, 2.5 µl of each reaction was resolved by SDS-PAGE on a 10% Tris-glycine gel and visualized by Coomassie blue staining. Quantitative PCR was subsequently performed to monitor the relative abundance of DHFR transcripts.

### RT-qPCR

To analyze the transcription level of DHRF in the in vitro translation reaction, 2 μl of the reaction mixture was treated with 10 U of RNase-free DNase Ⅰ (Thermo Fisher Scientific, MA, USA) to remove residual DNA, following the manufacturer’s protocol. The DNA-free product was then reverse transcribed using Moloney murine leukemia virus reverse transcriptase (M-MLV RT, Promega) to generate cDNA. For qPCR, 5 µL of a 50× diluted cDNA was used as template in a 20 µL reaction system prepared with the KAPA SYBR® FAST qPCR Kit (Kapa Biosystems). Amplification and detection were performed on an Applied Biosystems ViiA™ 7 Real-Time PCR System. Each sample was analyzed in triplicate. The primer sequences used for qPCR can be found in Table S2.

### Phage infection assay

The phage infection assay was performed as previously described with modifications^23^. Overnight bacterial culture (150 µL) was mixed with 3 mL of molten top agar (0.5% w/v) containing 10 mM MgSO₄ and corresponding antibiotics, and overlaid onto an LB agar (1.2% w/v) with identical supplements. After solidification, 2 µl of serial ten-fold phage dilutions were spotted onto the lawn. Plates were incubated overnight at 37℃. Following incubation, the plaques were counted, and the phage titer was calculated and expressed as Plaque-Forming Units per mL (PFU/mL). All experiments were performed with three biological replicates.

### CRISPR-Cas9 editing of the ΦYTSY4 phage genome

The genome of ΦYTSY4 was edited in *P. aeruginosa* PAO1 using a CRISPR-Cas9 counterselection strategy. All plasmids and oligonucleotides used in this study are listed in Table S2. First, a donor strain harboring a plasmid with a homology-directed repair template (flanked by 500-bp homology arms) was infected with the wild-type phage. The resulting lysate, containing a pool of phages enriched for homologous recombination events, was harvested. This lysate was then introduced into a selector strain expressing Cas9 and a target-specific sgRNA to selectively enrich for edited phages that evaded CRISPR cleavage. The progeny phage was subsequently passaged through fresh selector strains for three consecutive rounds. Finally, individual plaques were isolated, and the genomic modifications were confirmed by PCR amplification and Sanger sequencing.

### Plasmid conjugation

Plasmids constructed in this study are listed in Table S2. Derivatives of pRSF1010 and pVS1 were transferred into *P. aeruginosa* strains via conjugation using *E. coli* S17-1 λpir as the donor strain. Briefly, overnight cultures of the donor and recipient strains were mixed at a 1:2 ratio. The cell mixture was pelleted by centrifugation at 7,000 × g for 2 min and resuspended in 100 µL of LB broth. The concentrated cell suspension was spotted onto a sterile 0.45 µm filter membrane placed on an LB agar plate and incubated at 37 ℃ for 10 hours.

Following incubation, the conjugation spots were harvested and resuspended in 500 µL of LB. The resulting cell suspension was serially diluted 10-fold, and 200 µL of each dilution was plated on Pseudomonas Isolation Agar (PIA; BD Difco) containing appropriate antibiotics to select for transconjugants. To determine the recipient cell count, the initial mixture was also diluted and plated on non-selective media. Conjugation efficiency was calculated as the number of transconjugants (CFU/mL) divided by the number of recipient cells (CFU/mL). All experiments were performed with three biological replicates.

### Fluorescence measurement

A reporter construct was generated in which the *creT* promoter region (containing the CreA target sequence) drives the expression of the mCherry gene and was cloned into the pVS1 plasmid. The resulting reporter plasmid, along with a separate plasmid for CreA RNA expression, was then introduced into the recipient strain PAO1::Csy.

For measurement, transformants were inoculated into 3 mL of LB medium containing tetracycline, gentamicin, and carbenicillin and then grown for 10 hours until the OD_600_ reached approximately 0.8. Then, 200 µL of each culture was transferred to a microplate for the simultaneous measurement of OD_600_ and mCherry fluorescence (excitation/emission: 587/610 nm) using the Synergy H4 Hybrid multimode microplate reader (BioTek). For each of the three biological replicates, the ratio of fluorescence to OD600 was determined, and the mean and standard deviation were calculated. For statistical analysis, two-sided Student’s *t* tests were performed to determine significance.

### RNA isolation

Overnight cultures were diluted 1:100 into 10 mL of fresh LB medium containing the appropriate antibiotics and grown at 37 °C to the logarithmic phase. Bacterial cells from 2 mL of culture were harvested by centrifugation at 12,000 × g for 2 min at 4°C. Total RNA was subsequently extracted using the E.Z.N.A. Bacterial RNA Kit (Omega Bio-tek), following the manufacturer’s protocol. To eliminate genomic DNA contamination, the RNA samples were treated with DNase I (New England Biolabs) as instructed. RNA concentration and purity were assessed using a NanoDrop One Spectrophotometer (Thermo Fisher Scientific).

### Primer extension analysis

Primer extension was performed to map the transcriptional start site. A 5′-FAM-labeled primer specific to the *mCherry* transcript (Table S2) was used. For the reverse transcription reaction, 30 µg of total RNA was annealed with 2.5 µg of the labeled primer and extended with 200 U of M-MLV Reverse Transcriptase (Promega) according to the manufacturer’s protocol. The cDNA products were resolved on an ABI 3730xl DNA Analyzer, and the resulting data were analyzed with Peak Scanner software v1.0 to identify the transcription start site.

### Northern blot analysis

To analyze the CreA RNA products, the PAO141 or PAO142 strain was constructed by transforming PAO1 with a plasmid carrying the *creTA1* or *creTA2* module within the PA14 *cas* operon. For comparison, a control strain was generated by introducing an artificially designed mini-CRISPR system with a constitutively expressed Csy operon into PAO1.

To analyze the CreT RNA products, the plasmid carrying an L-arabinose-inducible *acrIF7* gene was introduced into the PAO141 or PAO142 strain. For induction, a single colony was grown to the logarithmic phase (OD600 approaching 0.6) in LB with antibiotics and then treated with L-arabinose for 3 hours. To analyze the effect of hairpin structures on the CreT RNA product, the wild-type *creT* gene and its mutant variants were cloned into L-arabinose-inducible plasmids and transformed into PAO1. Induction and RNA extraction were performed as described above for the *acrIF7* induction protocol.

For Northern blotting, a total of 10 µg of RNA was mixed with an equal volume of RNA Gel Loading Dye (Thermo Fisher Scientific), heated to 65 ℃ for 5 min and then cooled on ice for 2 min. Then, the RNA samples were resolved by denaturing 10% TBE-Urea PAGE and then electrotransferred onto Biodyne B nylon membranes (Pall). The RNA was electrotransferred onto a nylon membrane, UV-crosslinked, and pre-hybridized. Hybridization was performed overnight at 42°C with a biotin-labeled ssDNA probe. Signals were detected using a chemiluminescent kit (Thermo Fisher Scientific) per the manufacturer’s instructions and imaged with a Tanon 5200 system (Tanon Science & Technology). Each assay was repeated with separate biological samples, and a representative result is provided.

### Dilution plating assay

Individual colonies of *P. aeruginosa* were inoculated into LB broth containing the appropriate antibiotics and glucose (0.5% w/v), then cultured overnight to stationary phase. The cultures were subjected to 10-fold serial dilution in LB broth. A 2.0 μL aliquot of each dilution was spotted onto LB agar plates supplemented with the relevant antibiotics and either 1.0% (w/v) L-arabinose or 0.5% (w/v) glucose. After overnight incubation, colonies were counted to calculate the viable titer in colony-forming units per mL (CFU/mL).

### Animals and Ethics Statement

The Animal Research Ethics Committee of the Army Medical University reviewed, approved and supervised the protocols for animal research (permit number: AMUWEC20250084). BALB/c mice (7-week-old, female) used in this study were purchased from the Hangzhou Ziyuan Laboratory Animal Technology Co., Ltd. (Zhe jiang, China. License Number: SCXK (Zhe) 2024-0004). All the animals were housed under specific pathogen-free conditions; the housing environment had controlled temperature (20–26 °C), humidity (40–70%) and lighting conditions (12 h light and 12 h dark cycle), and no animal was excluded from the analyses.

### Mouse infection and phage therapy experiment

For the phage therapy experiment, 7-week-old BALB/c female mice were intraperitoneally inoculated with 100 μl of bacteria (PAO142 or PAO142m, ∼3 × 10^7^ CFUs). After 30min, 100 μl of phage ΦYTSY4, Δacr, ΦIF7, or ΦIF11 (∼3 × 10^8^ PFUs) was intraperitoneally inoculated into the infected mice. Each group included 5 mice, which were observed for 7 days. At 7 days after infection, mice that survived the initial challenge were euthanized.

## QUANTIFICATION AND STATISTICAL ANALYSIS

The number of replicates is specified in the associated figure legends. Each replicate represents a biological replicate of the specified experiment. Two-tailed Fisher’s exact test was performed for statistical analyses. *P*-values above 0.05 were considered non-significant. Statistical comparisons for the transformation assays relied on log values, which assumes the samples are normally distributed on a log scale.

## Supplementary Figures

**Figure S1.**
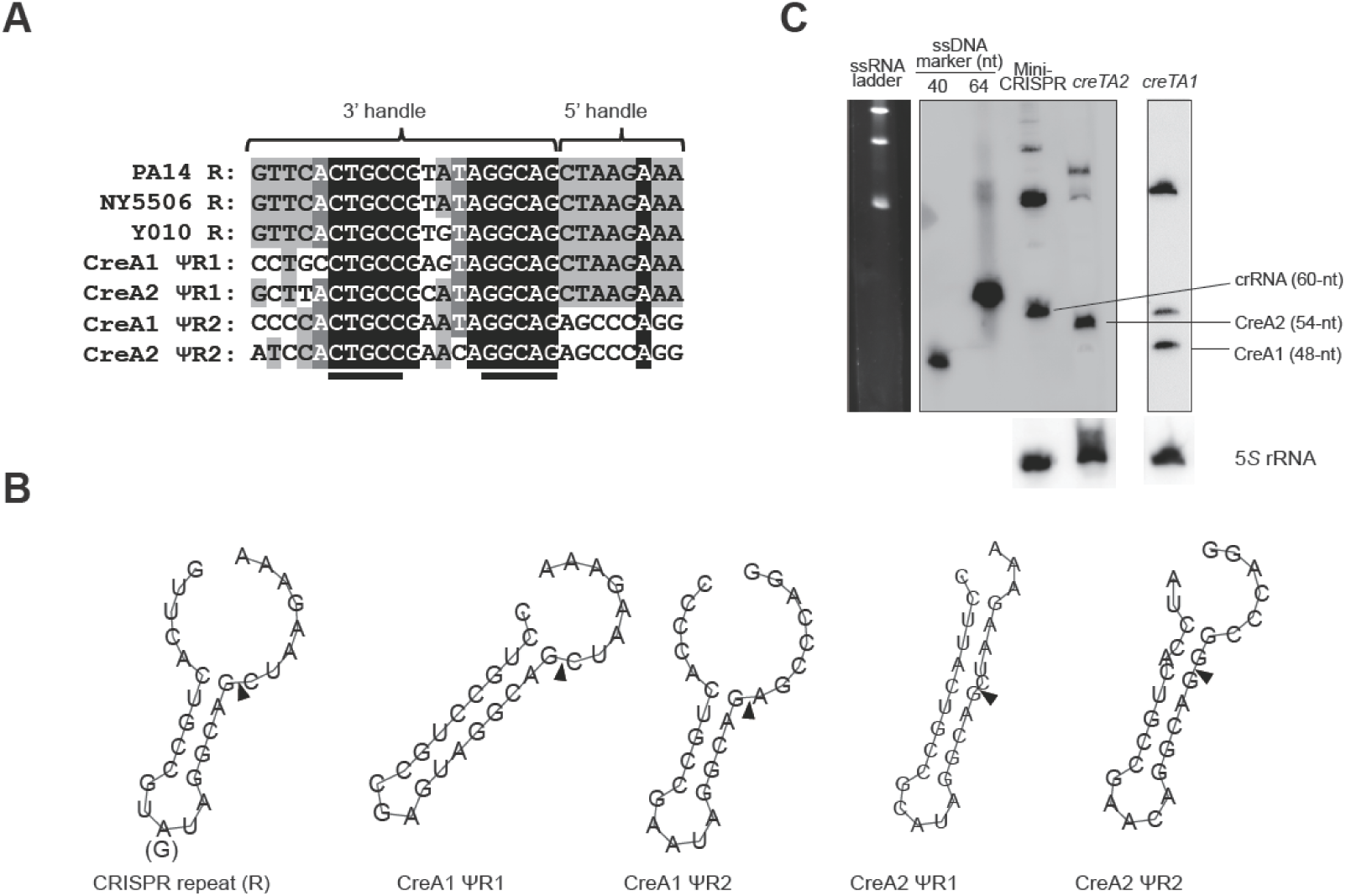
Hairpin-forming potential and Cas-processing of CreA repeat-like (ΨR) sequences, related to Figure 2. (A) Alignment of the ΨR sequences (ΨR1 and ΨR2) from *creA1* and *creA2* with the typical repeat (R) from the PA14, NY5506, and Y010 CRISPR arrays. Palindromic nucleotides are underlined. Nucleotides (nts) producing a 5’ or 3’ handle on mature RNAs are indicated. (B) Predicted structures of each repeat RNA. Triangles indicate the conserved processing sites by Csy4. (C) Northern blotting of CreA1, CreA2, and an engineered typical crRNA carrying the same spacer sequence as CreA2. All RNA samples were co-electrophoresed alongside a single-stranded RNA ladder and synthesized single-stranded DNA markers on the same gel, and following electrotransfer, the membrane was sectioned and hybridized with distinct probes. PAO1 expressing PA14 Cas proteins was used as the host.

**Figure S2.**
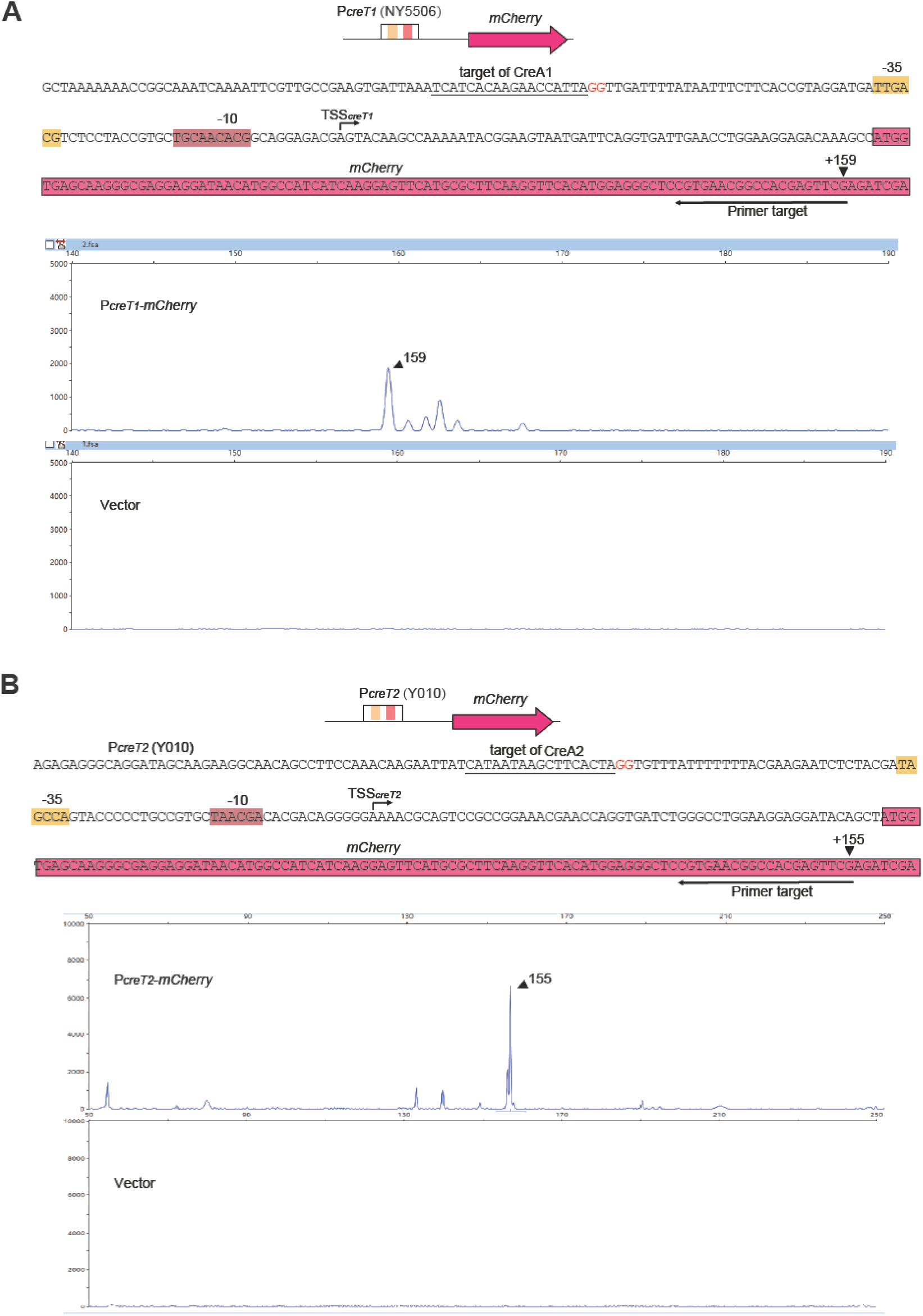
Characterization of the NY5506 P*_creT1_* (A) and the Y010 P*_creT2_* (B) via primer extension assays, related to Figure 2. (A) and (B) The promoter DNA was first linked to an *mCherry* gene and then introduced into *P. aeruginosa* PAO1 cells, of which total RNA was extracted for primer extension. The primer was designed against the *mCherry* RNA transcript and 5’-labeled with FAM (see the Methods section). The complementary DNA (cDNA) products from the primer extension assay were subjected to fragment size analysis. PAO1 cells containing an empty vector served as a negative control. The promoter elements −10 and −35 are indicated. The target sequence of CreA RNAs is underlined, with the complementary nucleotides of PAM shown in red.

**Figure S3.**
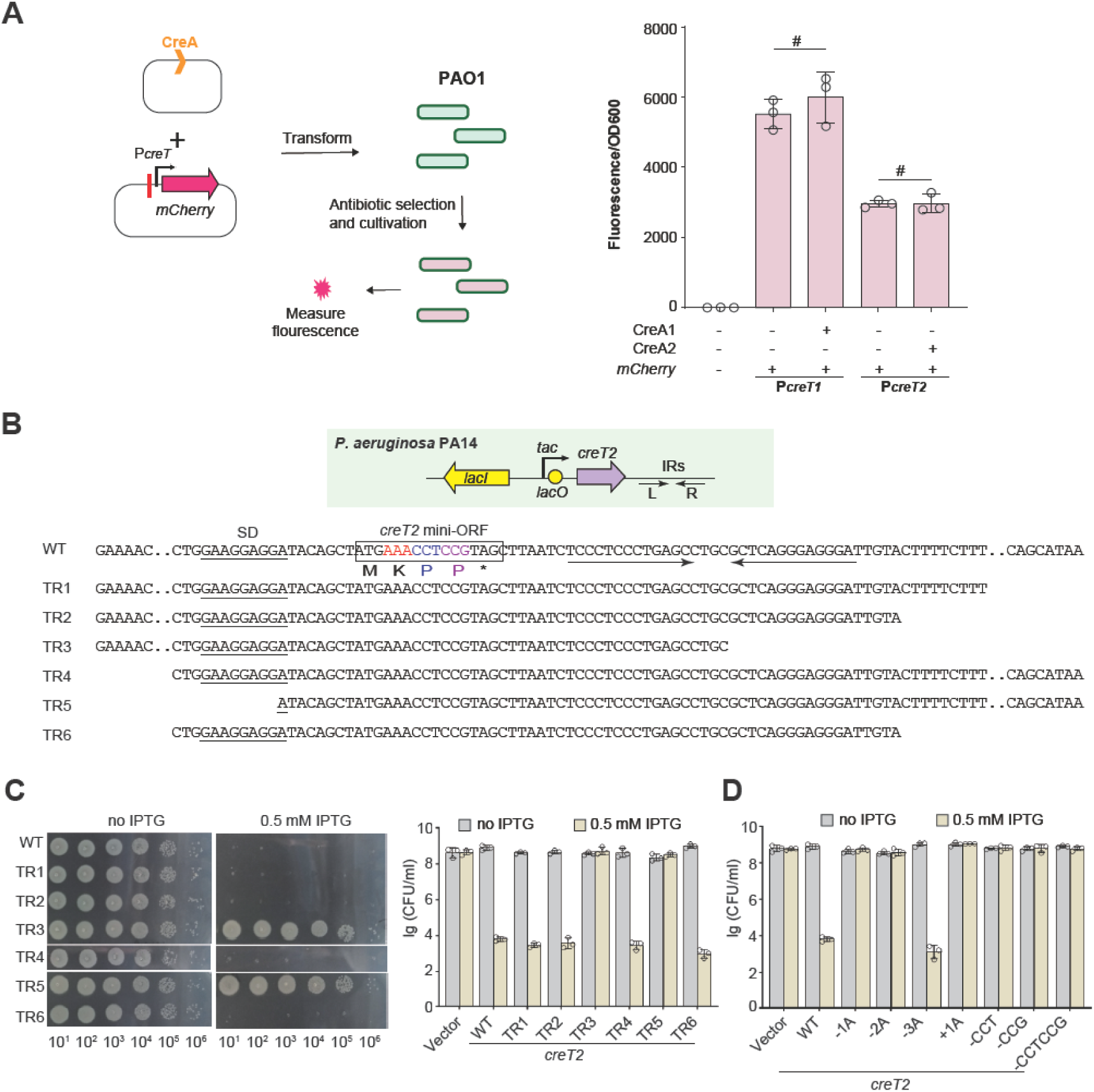
Truncation and frameshift mutation of the CreT2 RNA toxin, related to Figure 2. (A) Fluorescence intensity assay in the absence of Cas proteins. Error bars, mean±s.d. (n=3). *P* values were calculated using a two-sided Student’s *t* test. #, not significant. (B) Illustration showing the expression of differently truncated *creT2* sequences (TR1-TR6; under the control of *tac* promoter) to determine the minimal requirement for toxicity. The red nucleotides (AAA) were subjected to frameshift mutation analyses (panel **c**). The two proline codons (CCT and CCG) are highlighted in blue and purple, respectively. (C) Growth of PA14 cells expressing differently truncated *creT2* genes. (D) Cellular growth when the mini-ORF of *creT2* was differently mutated. One (−1A), two (−2A), or all (−3A) the three nucleotides of the AAA codon (shown red in panel **a**) were deleted for frameshift analysis. Addition of an extra adenine was also performed (+1A). The two proline codons were individually deleted in the -CCT and -CCG mutants, or simultaneously deleted in the -CCTCCG mutant. Error bars, mean±s.d. (n=3).

**Figure S4.**
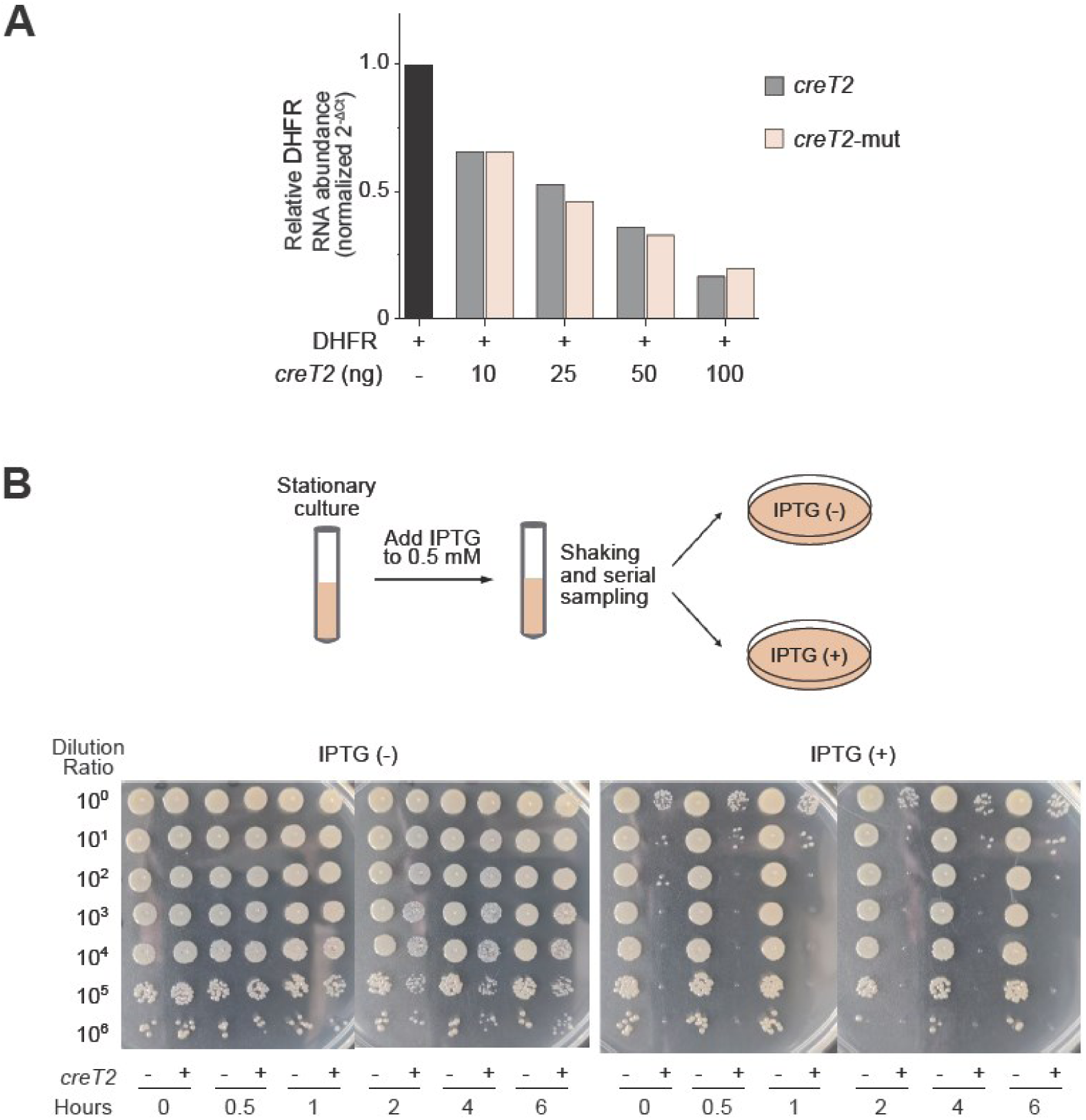
Characterization of CreT toxicity, related to Figure 2. (A) Quantitative PCR assay evaluating the effect of CreT2 on T7 polymerase-mediated transcription of the DHFR template. Values represent the mean of three technical replicates. In the *creT2-mut* control, the two critical proline codons were mutated. (B) Dilution plating assay. Stationary-phase PA14 cells carrying the IPTG-inducible creT2 construct were harvested at 0.5, 1, 2, 4, and 6 hours after IPTG addition, serially diluted, and plated on agar medium supplemented with or without IPTG.

**Figure S5.**
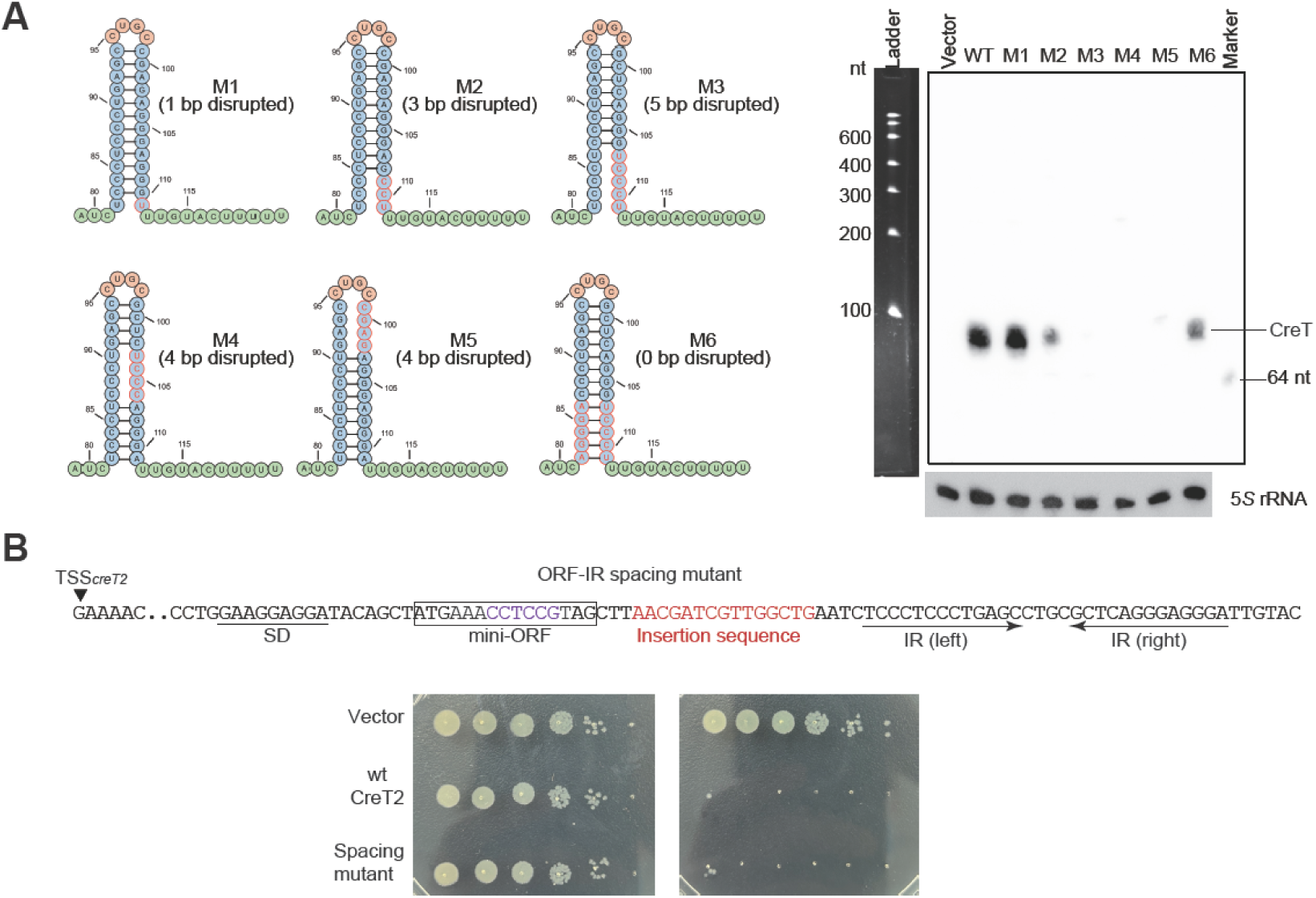
Mutational analysis of the CreT2 hairpin, related to Figure 2. (A) Northern blot of CreT2 mutants with a differently disrupted or restored hairpin structure. 5*S* rRNA served as the internal control. Biotin-labeled single-stranded oligos (40 nt or 64 nt) served as markers. A ssDNA ladder was run in parallel. (B) Toxicity of a CreT2 mutant with a 15-nt ‘spacing’ sequence inserted between mini-ORF and the hairpin.

**Figure S6.**
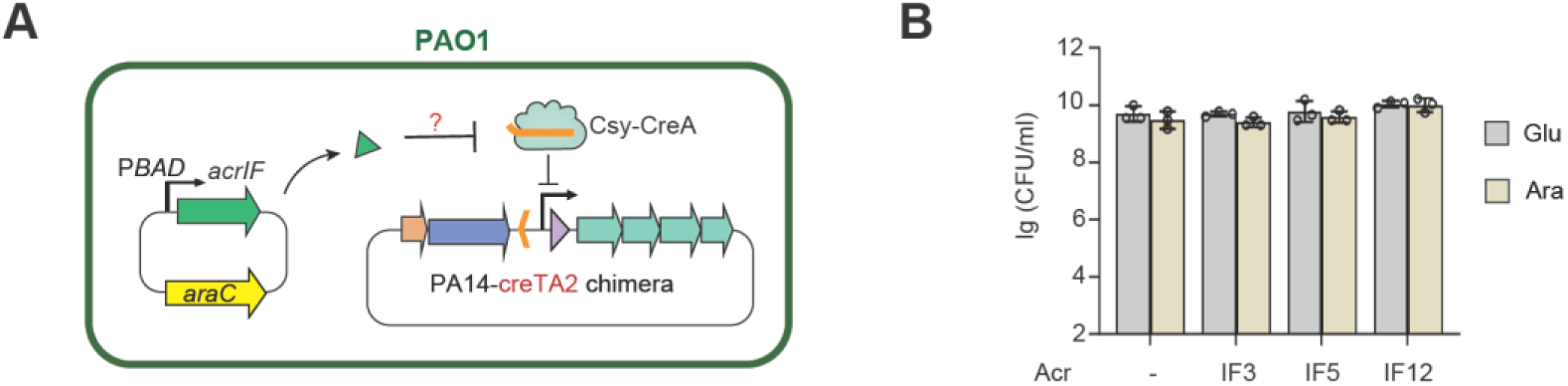
Activation of CreTA2 by Acr proteins, related to Figure 4. (A) Experimental design. The *acr* genes were placed under the arabinose-inducible PBAD promoter. The Y010 intergenic region encompassing *creTA2* was engineered into the PA14 *cas* operon to create a chimeric locus. (B) Growth of PAO1 carrying the PA14-CreTA2 chimera and an arabinose-inducible *acr* gene on medium supplemented with arabinose (Ara) or glucose (Glu). Error bars, mean±s.d. (n=3).

**Figure S7.**
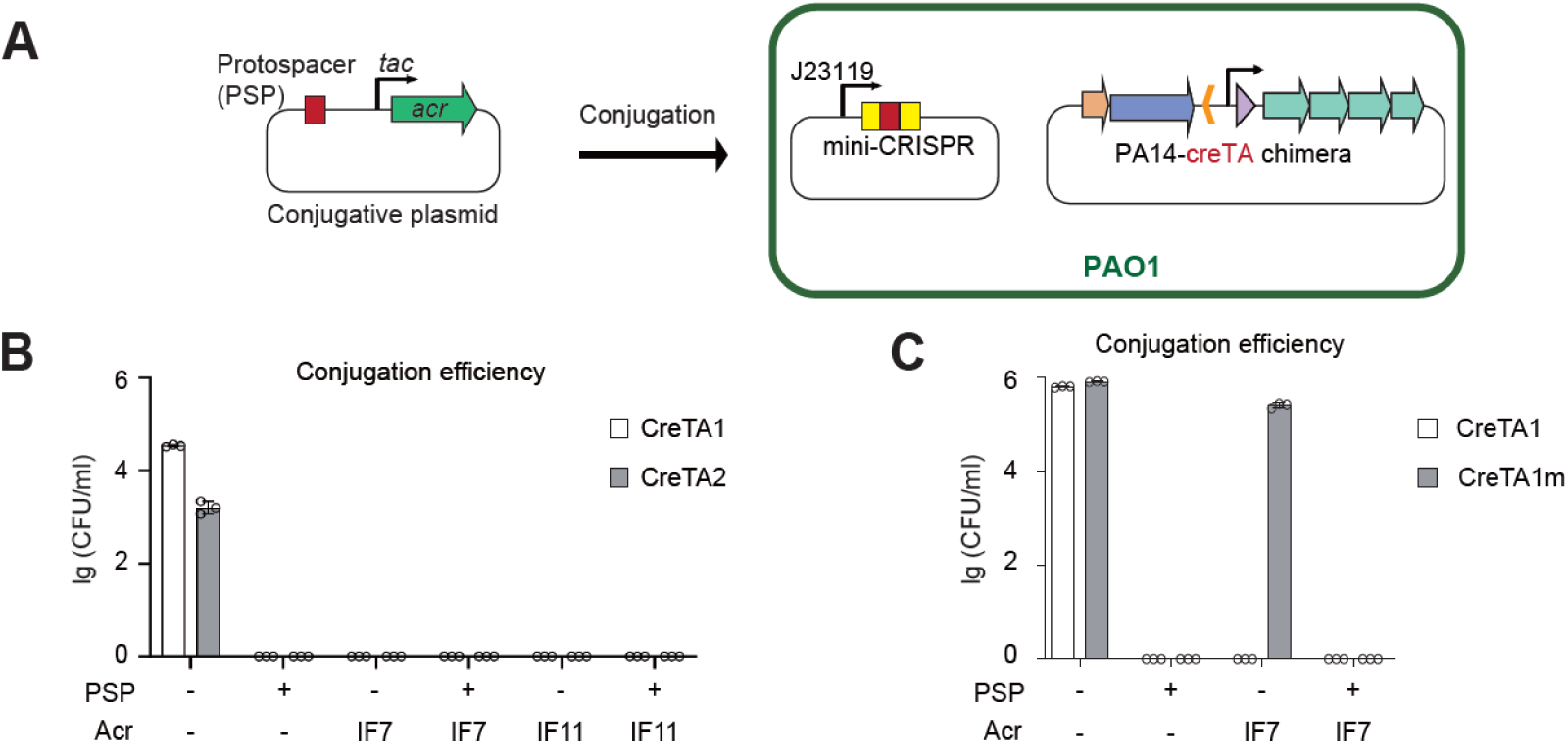
Conjugation of an *acr*-expressing plasmid into PAO1 cells harboring different CreTA modules, related to Figure 5. (A) Design of the conjugation assay. The host also carries a mini-CRISPR controlled by J23119. (B) Conjugation efficiency of PAO1 encoding wild-type CreTA1 or CreTA2. (C) Conjugation efficiency of PAO1 encoding wild-type or CreT-mutated CreTA1. ‘+’ indicates that the conjugative plasmid carries the cognate protospacer (PSP). Error bars, mean±s.d. (n=3).

**Figure S8.**
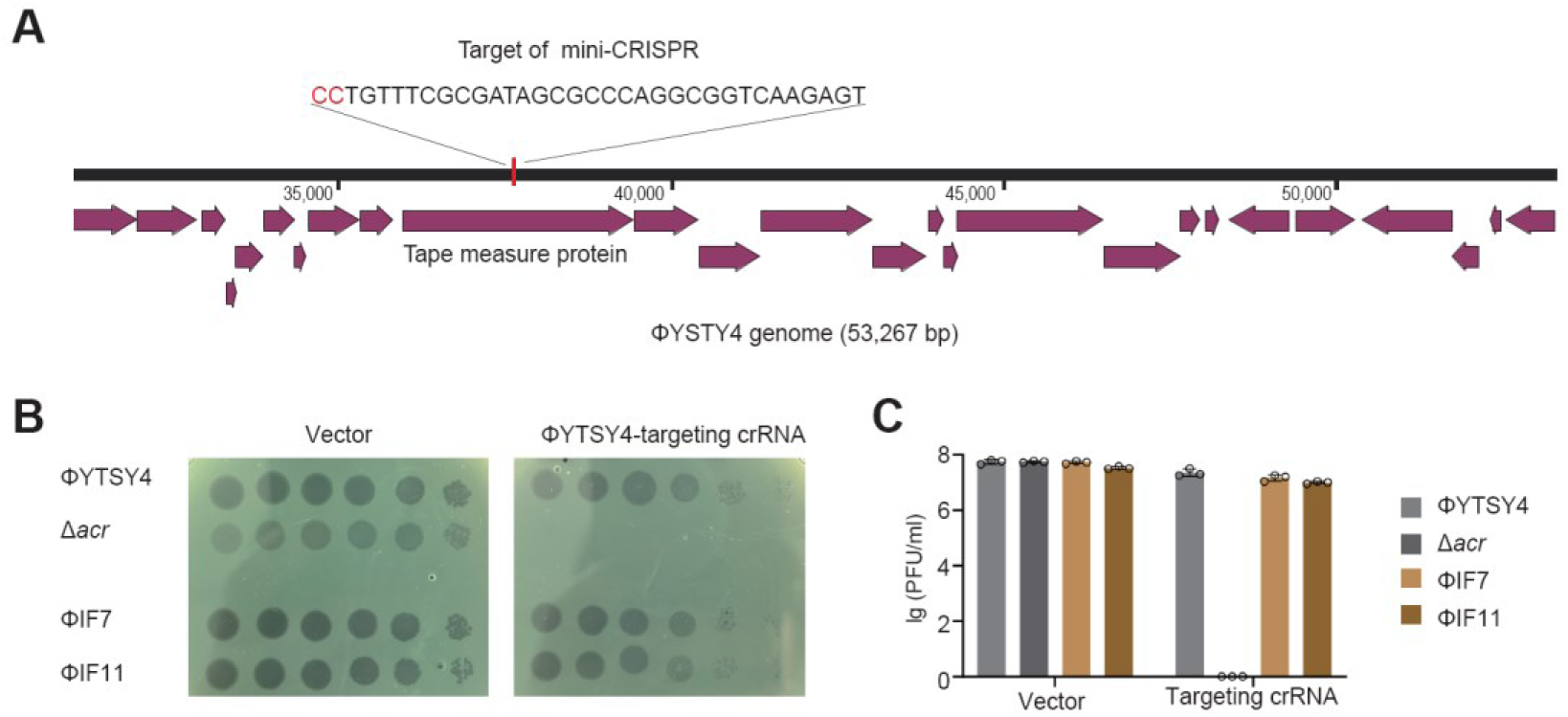
Escape of CRISPR-targeted immunity by ΦYTSY4 and its mutants, related to Figure 5. (A) The genome of ΦYTSY4 and experimental design. A mini-CRISPR was designed to target the gene of tail tape measure protein. The PAM is highlighted in red. (B) Plaques of serially-diluted phages formed on PA14 (with native CRISPR-Cas) cells expressing either phage-targeting crRNAs or an empty-vector control. (C) Statistical analysis of plaque forming units (PFUs). Error bars, mean±s.d. (n=3).

